# HELP: A computational framework for labelling and predicting human common and context-specific essential genes

**DOI:** 10.1101/2024.04.16.589691

**Authors:** Ilaria Granata, Lucia Maddalena, Mario Manzo, Mario Rosario Guarracino, Maurizio Giordano

## Abstract

Machine learning-based approaches are particularly suitable for identifying essential genes as they allow the generation of predictive models trained on features from multi-source data. Gene essentiality is neither binary nor static but determined by the context. The databases for essential gene annotation do not permit the personalisation of the context, and their update can be slower than the publication of new experimental data. We propose HELP (<u>H</u>uman Gene <u>E</u>ssentiality <u>L</u>abelling & <u>P</u>rediction), a computational framework for labelling and predicting essential genes. Its double scope allows for identifying genes based on dependency or not on experimental data. The effectiveness of the labelling method was demonstrated by comparing it with other approaches in overlapping the reference sets of essential gene annotations, where HELP demonstrated the best compromise between false and true positive rates. The gene attributes, including multi-omics and network embedding features, lead to high-performance prediction of essential genes while confirming the existence of essentiality nuances.

**Author summary:** Essential genes (EGs) are commonly defined as those required for an organism or cell’s growth and survival. The essentiality is strictly dependent on both environmental and genetic conditions, determining a difference between those considered common EGs (cEGs), essential in most of the contexts considered, and those essential specifically to one or few contexts (context-specific EGs, csEGs). In this paper, we present a library of tools and methodologies to address the identification and prediction of cEGs and csEGs. Furthermore, we attempt to experimentally explore the statement that essentiality is not a binary property by identifying, predicting and analysing an intermediate class between the Essential (E) and Not Essential (NE) genes. Among the multi-source data used to predict the EGs, we found the best attributes combination to capture the essentiality. We demonstrated that the additional class of genes we defined as “almost Essential” shows differences in these attributes from the E and NE genes. We believe that investigating the context-specificity and the dynamism of essentiality is particularly relevant to unravelling crucial insights into biological mechanisms and suggesting new candidates for precision medicine.

## Introduction

Identifying essential genes (EGs) is challenging and involves multiple disciplines and research areas. EGs are generally defined as necessary for the growth and survival of any organism or cell. The identification of essential genes was initially a prerogative of synthetic biology concerning the definition of the minimal genome [1]. In particular, EGs were considered those that cannot be removed or silenced from a genome without provoking a deleterious phenotype, reducing the organism’s viability or fitness. Technological advancement, on the one hand, and the clear potentialities emerging from EG research, on the other hand, led to the experimental scaling from microorganisms to more complex organisms, including humans. The accumulation of data and biological insights made the gene essentiality a key concept of genetics, with implications ranging from basic research to evolutionary, systems biology, and precision medicine [2]. Still, the definition is critical, as the term “essential" requires a specific contextualisation. In this scenario, a crucial role is played by the conditions in which the experimental procedures for the EGs recognition are performed. EGs are commonly identified through *in vitro* experiments on cell lines aimed at deleting the gene of interest and observing the effects on the phenotype. The more deleterious the phenotype, the more essential the gene. Single gene deletion, antisense RNA, transposon mutagenesis and CRISPR-Cas9 are the most used techniques. The latter is considered the state-of-the-art method for simplicity and efficiency [3]. At a genome-wide level, these procedures must be performed massively, becoming complex, costly, labour- and time-intensive. Computational methods to support the experimental approaches, minimising the costs and overcoming limitations related to the availability of *in vitro* models, are urgently needed. Approaches based on Machine Learning (ML) allow the generation of predictive models based on features coming from multi-modal, multi-source and multi-omics data, an aspect of primary importance in the era of precision medicine and the need for procedures able to capture context-specificity. The gene essentiality prediction is usually treated as a supervised classification problem, where the model is trained by using several characteristics of genes that are *a priori* labelled as Essential (E) or Not Essential (NE) [4, 5]. An efficient model requires both consistent data and labels. Along with the biological and genetic features that can capture the gene essentiality, a set of information can be derived from networks representing the interactions of biological factors. The information contained in these systems can be learnt unsupervised through Deep Learning (DL) techniques [6–8]. The network primarily used in the context of EGs prediction is the Protein-Protein Interaction (PPI) network, describing the physical connections among proteins. According to the centrality-lethality rule, the more central a gene, or its product, the higher its probability of being essential [9]. While the massive experiments on multiple cell lines and conditions evidenced that the essentiality of a gene is strongly dependent on the context and that gene essentiality is neither a binary nor a static property [10, 11], the computational approaches proposed seem to ignore these fundamental aspects, relying on a binary classification of organism-wide EGs. Besides those that are considered commonly essential (precisely defined common EGs), as essential to all or almost all contexts, and therefore involved in the vitality and reproduction of all the cells, some genes are essential only in specific contexts, where the context is given by the genetic and/or environmental background. In this scenario, the context can be meant as a tissue, a disease or a specific condition, and the genes are defined as context-specific EGs. The paper by Larrimore and Rancati [10] perfectly summarises this concept with a clear graphic explanation. Rancati et al. [11] give some examples of the potential therapeutic usage of this information for drug targeting and disease therapies. In a disease like cancer, where the cells reprogram themselves, the essentiality is expressed differently between healthy and disease conditions. Identifying and targeting these differentially essential genes would mean hitting the cancerous cells avoiding the healthy ones. Although the most investigated, cancer is not the only disease for which the individuation and characterisation of EGs is of great interest [12, 13]. A crucial issue concerning the prediction of EGs is the labelling process. In most cases, the annotation of human E genes is retrieved from dedicated databases. An important resource is the Online GEne Essentiality (OGEE) database [2], which provides both organism- and tissue-level data and the relative lists of EGs. The labels derive from gene knockout experiments that produce scores reflecting cell fitness in the wake of the deletion of a specific gene. However, a pre-compiled list represents a limitation for context-specificity where the context of interest can include or exclude some experiments. Furthermore, the update of these databases is often slower than the publishing of new experimental data. Beyond databases, some tools for identifying EGs based on gene deletion scores have been developed [14].

To address the aspects mentioned above, we present HELP (<u>H</u>uman Gene <u>E</u>ssentiality <u>L</u>abelling & <u>P</u>rediction), a computational framework for common and context-specific EGs prediction that treats both the labelling and classification tasks. HELP, schematically described in Fig 1, computes the labelling of genes as E/NE through an unsupervised approach. For the validation of the labelling algorithm applied to common EGs we exploited some reference sets (CENtools [15], Behan2019 [16], Hart2017 [17], and Sharma2020 [15]): we evaluated the overlaps of HELP-based cEGs labels with these sets. For the validation of HELP labelling applied to context-specific EGs, due to the lack of similar reference sets, we could only validate HELP-labelled csEGs by evaluating their overlaps with other labelling sets, such as those obtained by [14] and by OGEE in the contexts object of the study. For the development of csEG prediction models, we collected multi-source genetic features and the embedding extracted from human and tissue-specific PPI networks by a DL approach to investigate what characterises and can predict EGs. We individuated the best combination of gene attributes to capture the essentiality and demonstrated that integrating omics and network features improves the prediction performance. Finally, we exploited this flexible labelling method to explore the existence of essentiality shades.

**Fig 1.**
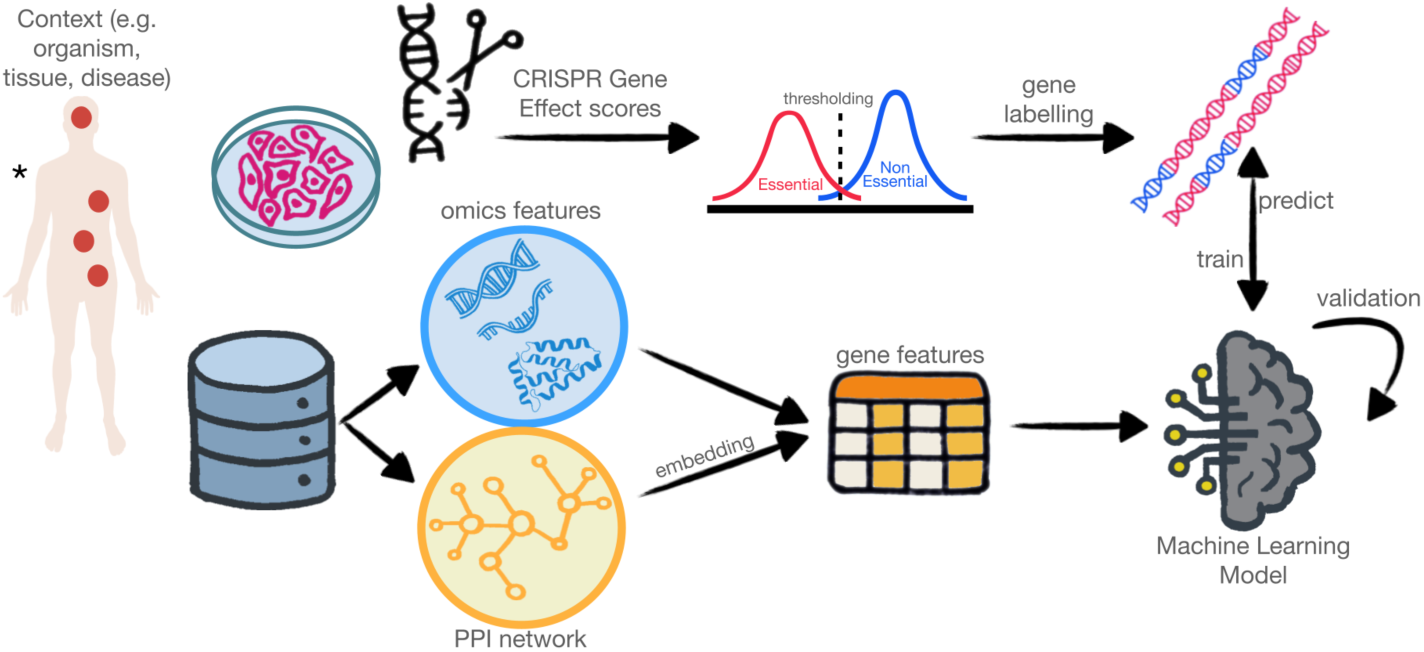
Schematic illustration of the HELP framework. HELP can be applied to each desired context (e.g. tissue, disease or the entire organism in the case of common EGs). The context guides the selection of gene knockout experiments and the collection of omics data and PPI. The CRISPR effect scores are used to derive the essentiality labels through an unsupervised thresholding approach. The omics and PPI embedding features are the input to train the machine learning prediction model. * Image from https://commons.wikimedia.org/wiki/File:Human_body_silhouette.svg.

## Materials and methods

### Labelling

The HELP framework labels genes based on the scores derived from gene knockout experiments. In particular, we used the scores reported in the Gene Effect file (DepMap v. 23Q4; https://depmap.org/portal), derived from the CRISPR knockout screens published by Broad’s Achilles and Sanger’s SCORE projects. As described in DepMap, negative scores imply inhibition of cell growth and/or death following gene knockout. Through the labelling step, HELP identifies common EGs (cEGs), context-specific EGs (csEGs) and uncommon context-specific EGs (ucsEGs), where the context (e.g. tissue or disease) is a user-defined parameter.

The methodology for identifying csEGs is the core of the labelling approach. For a chosen context, it consists in

1. selecting the knockout scores of the cell lines involved in the context,
2. for each cell line, automatically binarising the knockout scores to obtain E/NE cell line-dependent labels for each gene and
3. for each gene, assigning it the final E/NE label obtained as the mode of its cell line-dependent labels.

The selection in step 1 is obtained based on the annotations provided with the data, mapping each context to a subset of cell lines related to it. The binarisation in step 2 is obtained by thresholding the knockout scores, where the threshold is automatically determined by minimising intra-class intensity variance, or equivalently, by maximising inter-class variance, based on the Otsu method [18]; genes having scores lower/higher than the threshold are assigned an E/NE cell line-dependent label. For the computation of the mode in step 3, ambiguous cases (i.e. genes assigned an equal number of E/NE cell line-dependent labels) are solved by assigning NE as the final gene label.

Even though the HELP framework has been conceived focusing on context-specificity, it can also be adopted for identifying cEGs. To this end, our methodology consists of identifying csEGs for all the tissue contexts covered by the knockout data (using the above-described procedure) and assigning each gene the label obtained as the mode of its tissue-dependent labels. This strategy avoids the bias due to the different number of cell lines per tissue. Again, ambiguous cases in the mode computation are solved by assigning genes to the NE class.

Finally, as csEGs contain all the genes essential in the specific context, they implicitly include cEGs, which are therefore subtracted to obtain the very specific genes that we call ucsEGs.

In the experiments, we focused on two case studies, one concerning tissue-specific contexts and the other a disease-specific context, to extrapolate the differences between lung cancer subtypes. Moreover, as a pre-processing step, we filtered the CRISPR data by eliminating tissues having too few cell lines (*<*10) and genes having too many missing values across all cell lines (*>*95%).

### Prediction model

#### Training data

##### PPI embedding features

The human PPI network was downloaded from the STRING database v.13 [19]; we filtered out connections with a combined score below 0.5 to reduce false positive interactions. Tissue-specific PPIs were downloaded from the Integrated Interaction Database (IID) [20]. The edges of the IID human PPI are enriched with several annotations, including the tissue for which the interaction is reported and/or predicted. Filtering the PPI by tissue, we obtained Kidney-, Lung- and Brain-specific PPI networks of 19314 nodes/1110251 edges, 19334 nodes/1111550 edges and 19322 nodes/1122029 edges, respectively. Topological attributes of the PPIs were extracted using the node2vec algorithm [21]. It learns *node embeddings*, i.e. compact but informative vector-based representations of network nodes, by using the network topology as learning paths and maximising a neighbourhood-preserving objective function. The approach is flexible because the core of the embedding mechanism is a skipgram neural model [22] trained on simulated random walks and biased to allow different definitions of node neighbourhoods.

##### Multi-omics features

To achieve the prediction task and investigate the characteristics descriptive of essentiality, we collected generic and context-specific genetic attributes from several sources. We generated Kidney (3331), Lung (3330) and Brain (3330) numeric features regarding genomic, transcriptomic, epigenetic, functional, evolutionary and disease-related characteristics, summarised and detailed in Table S.1. For the organism-wide context (from now on referred to as Human) and prediction of cEGs the multi-omics features were 3323.

Structural information was represented by attributes such as “Gene length", “GC content", and “Transcript counts".

Four gene expression-related attributes were considered: “GTEX_*" (*= Kidney, Lung, Brain or Human), median-normalised gene expression values in the tissue (for Kidney, both medulla and cortex have been considered; for Human the median over all the tissues has been calculated) from GTEX portal [23]; “UP_tissue", the number of tissues in which the query gene has been found expressed as annotated in DAVID [24]; “OncoDB_expression", mean of the *log*_2_Fold-Change values as results of the differential expression analysis of three renal cancer (KIRC, KIRP, KIRC), two lung cancer (LUAD, LUSC) and one brain cancer (GBM) molecular subtypes vs normal samples [25]. Only genes with adjusted p-value *≤*0.05 were considered; “HPA_*” (*= Kidney, Lung, Brain or Human), normalised transcript expression values summarised by genes from RNA-sequencing experiments (for Human the median over all the tissues has been calculated) [26].

Functional characteristics were retrieved from Gene Ontology (GO), KEGG and REACTOME annotations. To convert the textual annotations into numerical attributes, for each query gene, we calculated the number of GO-Molecular Functions (“GO-MF”), GO-Biological Processes (“GO-BP”), GO-Cellular Components (“GO-CC”), KEGG (“KEGG”) and REACTOME (“REACTOME”) annotations. Attributes concerning protein sub-cellular localisation were also collected as confidence scores assigned to each of the 3305 GO-CC terms retrieved (“CCcfs”) from COMPARTMENTS [27]. Each term constitutes a single attribute.

As EGs are genes strongly involved in physical and functional interactions, we considered two interactions-based attributes: “BIOGRID”, the number of interactions annotated for each query gene [28], and “UCSC_TFBS”, the count of predicted transcription factors binding sites (TFBSs) from the UCSC TFBS.

EGs and their functions are reported to be highly conserved. We added the number of orthologs for each query gene as a measure of conservation (“Orthologs count”), retrieved from [29].

In the context of cancer, EGs strongly overlap with cancer-driver genes, in which genetic mutations determining uncontrolled cellular growth occur with the highest frequency. For this reason, we collected cancer-specific data from DriverDBv3, which provides three types of driver sets according to the kind of alteration identified through multiple dedicated bioinformatics algorithms [30]. We counted, for each query gene, the times in which it is predicted as a Mutation driver (“Driver_genes_MUT”), a Copy Number Variation driver (“Driver_genes_CNV”) or a Methylation driver (“Driver_genes_MET”) in all cancers and then specifically for renal (KIRCH, KIRP, KICH), lung (LUAD, LUSC) and brain (GBM, LGG) cancer subtypes. To characterise the genes according to their association with human diseases beyond cancer, we added the attribute “Gene-Disease association”, in which the counts of these associations are reported as annotated in DisGeNet [31].

##### Features’ sets

The features described in the previous paragraph and those extracted through graph embedding were organised into three sets to be evaluated in the downstream experiments:

###### Bio

Kidney - 28 attributes, Lung*Î*Brain - 25 attributes, Human - 18 attributes (consisting of all the features described in Table S.1 except “CCcfs”);

###### CCcfs

3305 attributes (see Function and Localisation features in Table S.1);

###### N2V

128 attributes, consisting of the node embeddings extracted as described in Section *PPI embedding features*.

##### Feature data pre-processing

Gene feature data were pre-processed by removing attributes having constant values. In the case of the Kidney context, we found two attributes with constant values in the Bio set (“Driver_genes_CNV (KICH)”, “Driver_genes_MET (KICH)”), which are context-specific and related to tumour subtypes. In the case of the Lung context, instead, none of the Bio attributes was constant, while for Brain two attributes were removed as constant (“Driver_genes_MET_GBM”, “Driver_genes_MET_LGG”). For Human, three attributes with constant values were found in the CCcfs set (“Integral component of endoplasmic reticulum-Golgi intermediate compartment (ERGIC) membrane”, “Intrinsic component of endoplasmic reticulum-Golgi intermediate compartment (ERGIC) membrane”, “Pentameric IgM immunoglobulin complex”). In addition, before the prediction model training, all feature sets except the N2V set were normalised using the z-score technique.

#### The meta-model classifier

We designed a new machine learning method [32], named <u>S</u>plitting <u>V</u>oting <u>E</u>nsemble (SVE), consisting of a soft-voting ensemble of *n* classifiers exploiting a splitting strategy on the training dataset based on labelled samples distribution (see Fig 2). The proposed method can be considered a *meta-learning algorithm* since it uses another learning method as a base model for all members of the ensemble to combine their predictions. This algorithm was designed and developed to address the strong class unbalancing inherent to the EGs prediction, although its applicability goes beyond the specific domain and can be extended generically to improve the classification performance in highly unbalanced datasets. In the following, we describe the logic and algorithmic operation of the proposed method. Before training, the method partitions the set of majority class samples into *n* parts, and it trains each classifier on a subset of training data composed of one of these parts along with the entire set of minority class training samples (Fig 2A). During testing on unseen data, each classifier of the ensemble produces a probability for the label prediction; we compute the final probability response of the ensemble as the average of the probabilities of the *n* voting classifiers (Fig 2B). The number *n* of classifiers is chosen based on the class data distribution of training data to train each of the *n* classifiers on a balanced subset of data.

**Fig 2.**
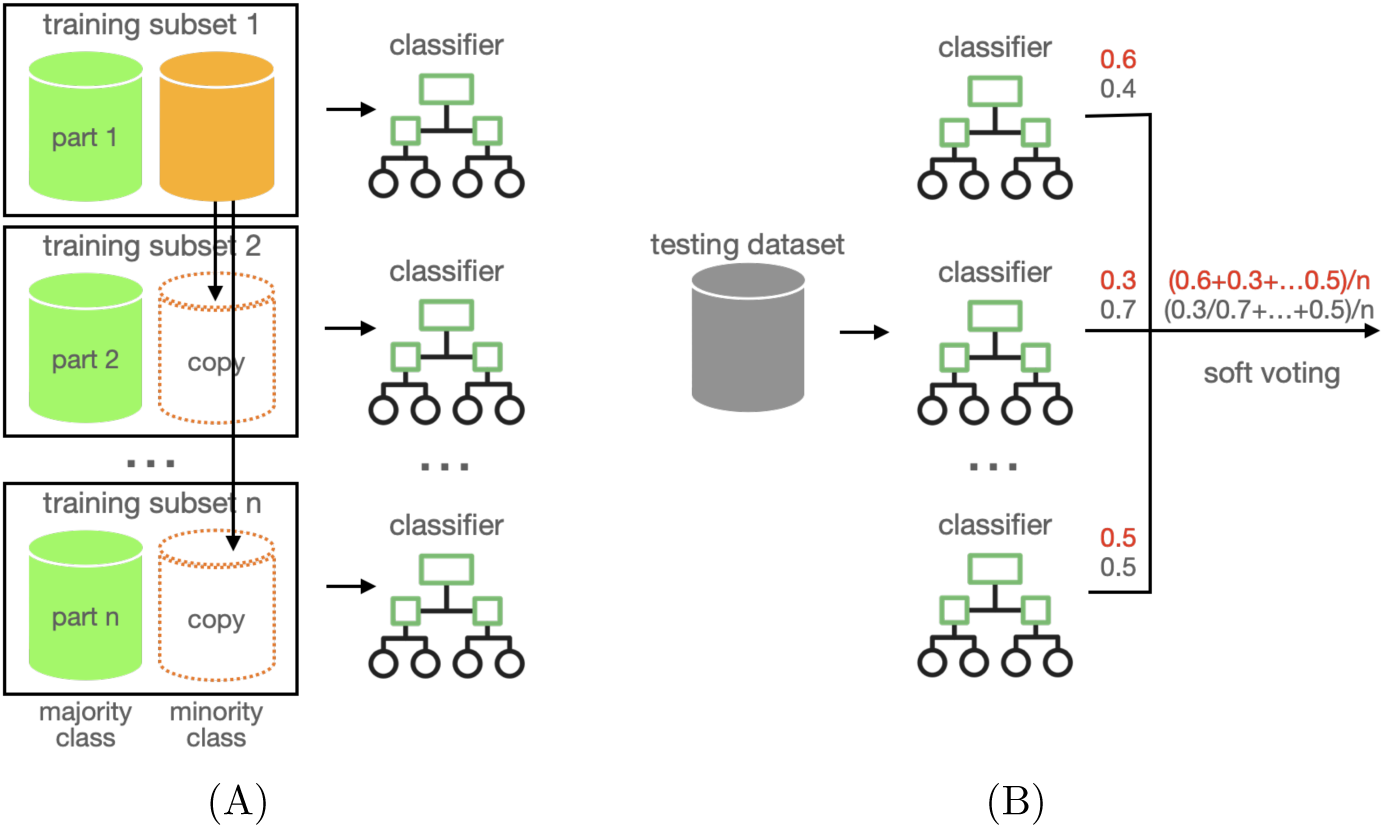
Splitting voting ensemble of classifiers: (A) In the training stage, the majority class samples are partitioned into *n* slices, each one coupled with the minority class samples to form datasets as inputs for the *n* trained models. (B) In the testing stage, each new sample is predicted by each of the *n* classifiers, and the final probability response is the average over the *n* partial responses.

For the current study, we first conducted preliminary tests to select the optimal base model of our ensemble (see section M.1 in Supplementary Material document and Table S.2). By using our method with a base classifier like LGBM, Random Forests [33] or AdaBoost [34], we outperform existing models in the PyCaret library in terms of BA and Sensitivity. The top-ranked method is our splitting voting ensemble of LGBM classifiers [35]. These experiments enforce our ensemble design of estimator acting on a split of the original data, which encompasses strong unbalancing of labelled samples. The counterpart of choosing sveLGBM is time costs: we pay a timing overhead in executing an ensemble of LGBM classifiers, each one of them being by itself ensembles of decision trees. In our opinion, this cost is reasonable for the gain obtained in performance. Consequentially to these results, we named sveLGBM the implementation of our meta-model as an ensemble of LGBM classifiers used in the current work. The hyper-parameter configurations for each classification problem were defined through optimisation tests (see section M.2 in Supplementary Material document and Table S.3) and the results discussed in section *Validation of cEG and csEG prediction*.

## Results

The purpose of the experimental study we conducted was threefold: 1) to demonstrate the effectiveness of HELP labelling in identifying common EGs (cEGs), context-specific EGs (csEGs) and uncommon context-specific EGs (ucsEGs); (see sections *Validation of common EGs (cEG) identification* and *Validation of context-specific EGs (csEG) and uncommon csEGs (ucsEG) identification*); 2) to develop and validate a supervised classifier to predict cEGs and csEGs on the basis of an optimal set of gene attributes representative of gene essentiality (see section *Validation of cEG and csEG prediction*); 3) to investigate the hypothesis of a class of genes that show an intermediate behaviour between E and NE, providing proofs through the computational approach and biological interpretation of the results (see section *Investigating almost essentiality*).

### Validation of common EGs (cEG) identification

The HELP framework has been thought and created for identifying and predicting specifically the csEGs, even if it also allows the identification of the cEGs. To prove the effectiveness of the unsupervised HELP labelling, we intersected the cEGs computed as described in Section *Labelling* with some sets used as reference in [14] (Fig 3A). As the diagram shows, the overlap among all the sets contains 324 genes. The HELP labelling agrees with CENtools [15] for 970 genes, Behan2019 [16] for 491, Hart2017 [17] for 584, and Sharma2020 [15] for 999 cEGs. The best overlap was therefore achieved in the case of the more recent sets of cEGs, i.e. CENtools and Sharma2020.

**Fig 3.**
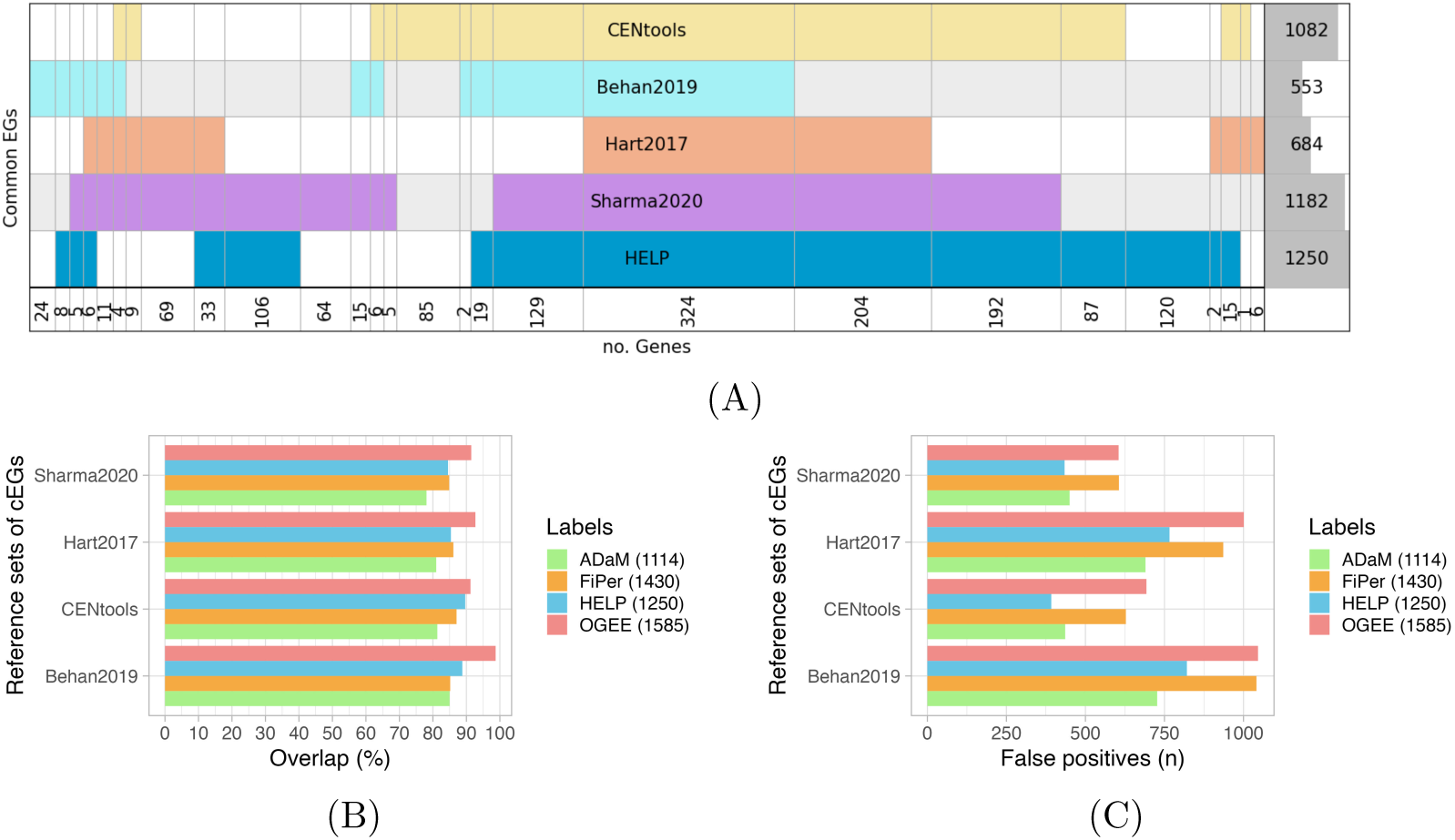
Comparison of HELP labelling with reference sets of cEGs. (A) Diagram representing the intersection of E genes labelled by HELP with reference sets of cEGs. Each row represents a different set of E genes. The last row reports the number of genes that result from the intersections. The last column on the right indicates the number of E genes for each set, with the dark grey shadow representing the corresponding histogram. (B) ADaM, FiPer, HELP and OGEE cEGs overlap with the reference sets of cEGs. In the legend, the number of cEGs identified by each approach is reported in brackets. (C) ADaM, FiPer, HELP and OGEE cEGs false positives, namely the genes not overlapping with the reference sets of cEGs. In the legend, the number of cEGs identified by each approach is reported in brackets.

To further demonstrate the reliability of cEGs identification by HELP, we compared it with the approaches proposed by [14], implemented into the CoRe R package, and the list of cEGs provided by the OGEE database. The two methods of the CoRe package, the supervised Adaptive Daisy Model (ADaM) and the unsupervised Fitness Percentile (FiPer), were used on the CRISPR data matrix filtered on columns (cell lines) and rows (genes) as explained above. In the case of ADaM, as the method requires a binarised matrix, we first converted the input to binary scores by setting the threshold suggested in the package documentation (*≤≠*0.5= *E*), then used the function “CoRe.PanCancer_ADaM”. In the case of FiPer, we derived a consensus set of predicted E genes by intersecting the output of the three less stringent FiPer variants (Fixed, ROC-AUC, Slope). The comparison was performed through the evaluation of the overlaps with the reference sets of cEGs mentioned above (Fig 3B). The OGEE list provided the greatest overlap with all four sets. OGEE considers common genes showing essentiality in at least 50% of cell lines. Such a low threshold determines a high number of cEGS (1585), thus including many EGs of the reference sets but with a high number of false positives (Fig 3C). The results by FiPer and ADaM confirmed what was commented by their authors regarding the stringency of the two methods (1114 ADaM, 1430 FiPer). The stringency grade influences the risk of erroneously identifying EGs. Indeed, FiPer detected much fewer cEGs than OGEE but with comparable false positives. Adam, instead, showed the best results in terms of false positives but with very low overlap percentages. In this scenario, HELP seems to provide the best compromise between the two evaluations. In two cases (Sharma2020, CENtools), it returned the lowest number of false positives and was the second-best in terms of overlap percentages.

### Validation of context-specific EGs (csEG) and uncommon csEGs (ucsEG) identification

To demonstrate the effectiveness and usefulness of HELP, we applied it to two different case studies: tissue- and disease-specific EGs.

#### Case study 1: tissue-specific EGs

We tested HELP in three tissue contexts: Lung, Kidney and Brain. The distributions obtained for the classes E/NE in the three contexts are reported in Fig 4A-4C. The E genes in all three cases are almost 7%, in accordance with the 10% reported in the literature.

**Fig 4.**
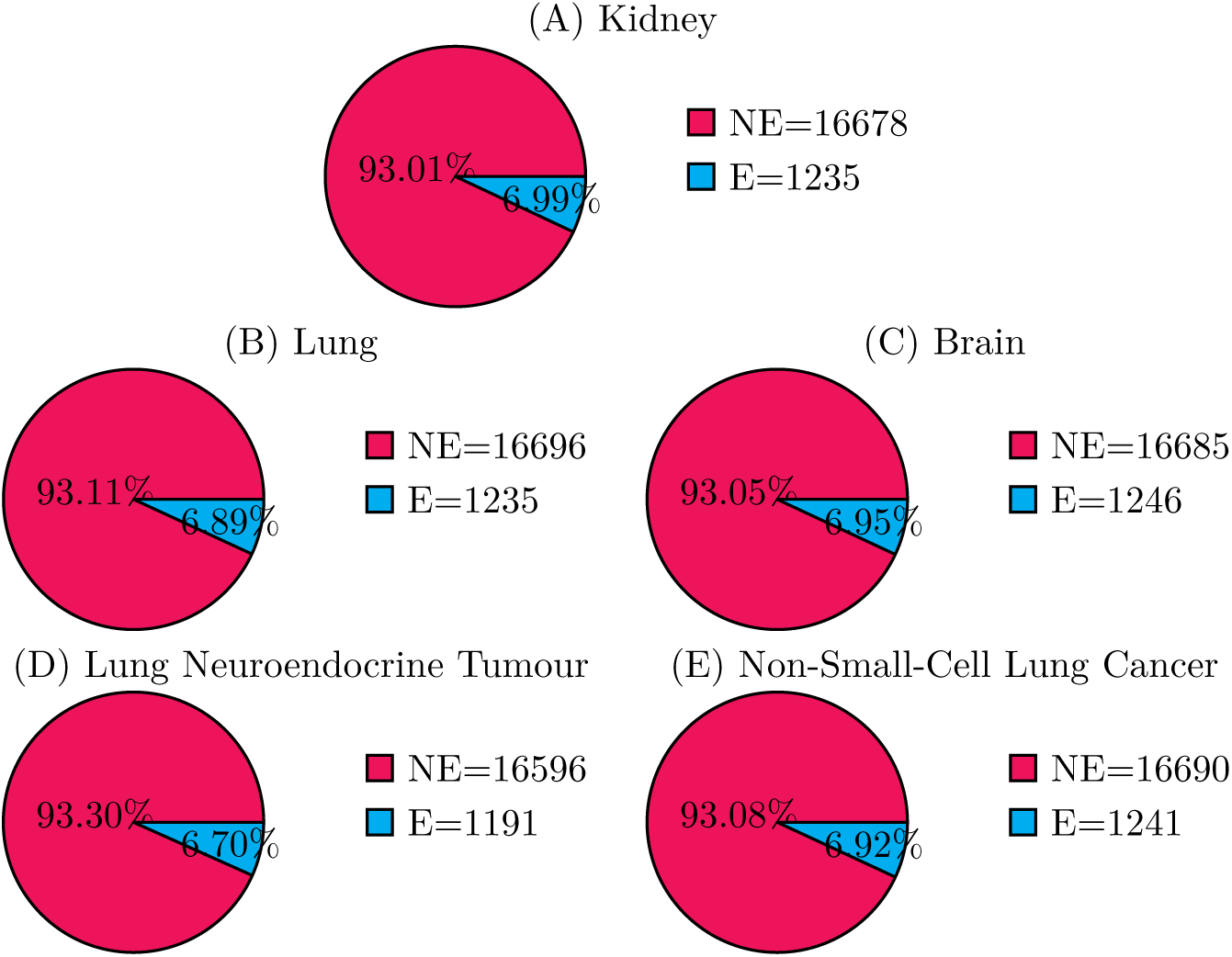
Distribution of the csEGs according to HELP labelling. (A) Kidney, (B) Lung, and (C) Brain tissues; (D) Lung Neuroendocrine Tumour and (E) Non-Small-Cell Lung Cancer. The count (in the legend) and percentage (inside the pies) of labelled genes are reported.

Although a standard validation is not possible due to the lack of experimentally validated context-specific EGs, to evaluate the ability to identify the csEGs, we compared the obtained results with tissue-specific EGs identified by the CoRe’s methods and those annotated in the OGEE database. To apply the ADaM method by CoRe, we used the “CoRe.CS_ADaM" function after binarising the input matrix, while in the case of FiPer, we just selected the cell lines of the tissue of interest from the score matrix and considered the E genes from the consensus set (Fixed, ROC-AUC, Slope). Looking at the number of csEGs obtained (Fig. 5), the most stringent sets are those provided by OGEE, even if, as already observed for cEGs, the threshold on cell lines for considering a gene essential in a tissue is 50%. The higher stringency of ADaM compared to FiPer is not confirmed in the case of csEGs. The intersection sets between all the csEGs included 742, 830 and 993 genes for Kidney, Lung and Brain, respectively. The highest overlaps are between HELP and ADaM in the case of Kidney (1192 genes) and between ADaM and FiPer in the case of Lung and Brain (1271 and 1292, respectively). In the Kidney and Brain contexts, FiPer showed the highest number of genes not shared with the others (79 and 76, respectively), while for Lung, ADaM provided the largest set of "unique" genes (200).

**Fig 5.**
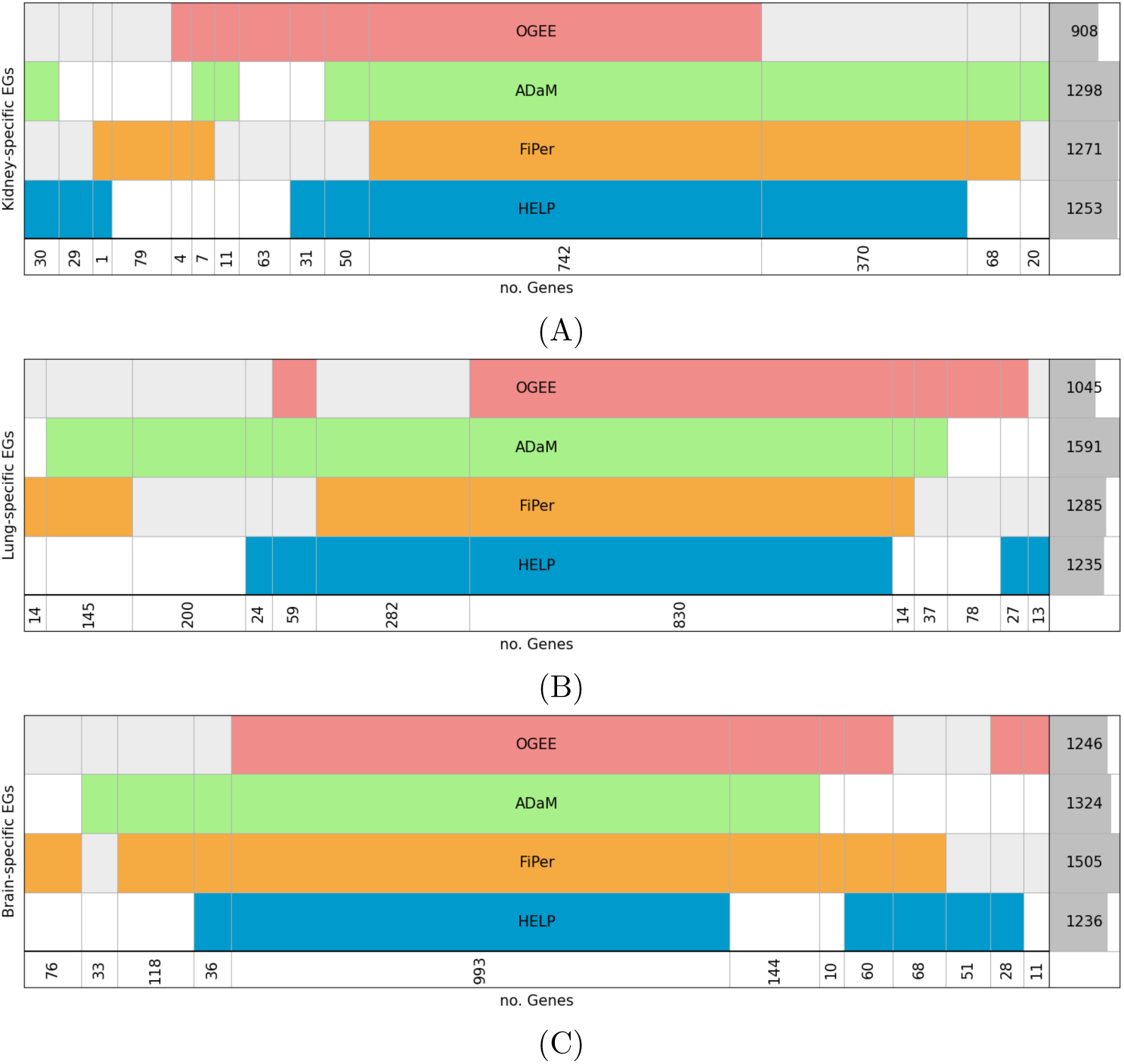
Comparison of HELP labelling with CoRe’s and OGEE csEGs. (A) Kidney, (B) Lung and (C) Brain tissue. Diagram representing E genes intersections among ADaM, FiPer, OGEE and HELP labelling. Each row represents the set of E genes for a labelling. The last row reports the number of genes resulting from the intersections. The last column on the right indicates the number of E genes for each set, with the dark grey shadow representing the corresponding histogram.

In our opinion, we cannot really talk about csEGs if we do not exclude from their computation the genes identified as cEGs, i.e. considering what we named ucsEGs. In the three tissues considered in this work, using the HELP framework, we obtained 28, 60 and 42 ucsEGs for Lung, Kidney and Brain, respectively. The difference in numerosity could be ascribed partially to the different number of cell lines in the input data, making the results of the context with more data (i.e. Lung) more stringent, but likely also to the different nature of the tissues, where kidney and brain are histologically more heterogeneous. The enrichment analysis of the three sets (Fig S.1) highlighted that, besides the context-specificity, the EGs maintain their identity characteristics, such as the main localisation in intracellular organelles, e.g. nucleus and mitochondrion, (GO Component “Intracellular membrane bounded organelle", “Intracellular organelle lumen"), or their involvement in growth functions (e.g. “Signal Transduction", “Cell cycle", “Cellular component biogenesis"). Most of them (41/60 in the Kidney and 22/28 in the Lung) contain the keyword “phosphoprotein" in their UniProt description, where protein phosphorylation is the key process to transmit extracellular and intracellular signals. Two genes were shared among the three tissues. Kidney and Brain showed the highest sharing, with seventeen genes in common. The results suggested that, besides the subtraction of the cEGs, some genes can be essential in different contexts and, then, rarely unique to a single one.

#### Case study 2: disease-specific EGs

To demonstrate the potential use of HELP as a tool for identifying candidate disease biomarkers, we applied HELP labelling to individuate disease-specific EGs in two lung cancer subtypes, Non-Small-Cell Lung Cancer (NSCLC) and Lung Neuroendocrine Tumour (NET), that account for 85% and 15% of lung cancer, respectively. The distributions of the classes E/NE in the two contexts are reported in Fig 4D-4E. The labelling was performed using the same procedures described for the tissue contexts, and the results were evaluated by comparing them with CoRe’s methods. The NET- and NSCLC-specific EGs were obtained by subtracting the common EGs and intersecting the ucsEGs of the two subtypes and those of the Lung (Fig S.2). The cEGs subtracted werecalculated by the mode of the tissue-specific labels. The idea is that the list of cEGs, namely the organism-wide EGs, must be unique. In our approach, the different tissues with different cell populations were considered as they are what make up the human organism. The three methods gave rise to different numbers of NET- and NSCLC-specific EGs (8, 6 and 23 NSCLC-specific EGs, 43, 22 and 97 NET-specific EGs by HELP, ADaM and FiPer, respectively). FiPer was the less stringent, although the number of ucsEGs before the intersection between the two diseases was not the highest (last column of the diagram in Fig S.2); on the contrary, HELP was the most stringent both before and after the intersection. The NET-specific EGs identified by HELP enriched pathways related to cancer cells’ progression and drug resistance strategies involving cellular respiration and energy production (Fig S.3) and that are particularly stressed in NETs [36]. The eight NSCLC-specific EGs were too few to perform an enrichment analysis. Still, we observed that they are all significantly differentially expressed when comparing the two NSCLC subtypes (LUAD and LUSC) with the normal samples (Fig S.4).

### Validation of cEG and csEG prediction

The prediction of cEGs and csEGs has been validated by 5-fold stratified cross-validation. Each cross-validation round, with a different random partition of the validation folds, was iterated ten times. In each validation step of the iteration, all genes in the dataset were predicted in such a way that we were able to evaluate means and standard deviations of single gene predictions as well as of the collected performance metrics (described in Table S.4).

#### Features set for predicting EGs

We reported the prediction results (Fig 6 and Table S.5) in terms of several metrics for evaluating the classification performance to allow easy comparisons, although aware that the most reliable and informative one is the Balanced Accuracy (BA), which in binary classification is the arithmetic mean of Sensitivity and Specificity (Table S.4), recommended for its superior robustness to imbalance and its applicability to both binary and multi-class prediction [37]. The evaluation of each set alone showed that we obtained adequate results in all the cases. The highest performance was in charge of CCcfs and the lowest of Bio attributes. Despite this, the Bio set is the most varying one in terms of gene characteristics and contains the context-specific attributes necessary for the prediction of ucsEGs. By combining the CCcfs and Bio attributes, we noticeably improved the performance. The N2V set containing the PPI networks’ embedding-derived topological attributes is also a context-specific attribute and alone gave a good performance. Its addition to the Bio+CCcfs set determined a slight improvement in the performance for all the tissue cases (Kidney - BA=0.892±0.009; Lung - BA=0.895±0.009; Brain - BA=0.895±0.008), a modest improvement Sensitivity (Kidney: 0.878±0.005; Lung: 0.879±0.005; Brain: 0.881±0.007), and a higher improvement in Specificity (Kidney: 0.905±0.019; Lung: 0.910±0.018; Brain: 0.910±0.018). This result confirmed the contribution of N2V attributes specifically to the context-specific prediction. We then decided to consider the combination including all the attributes as the best-performing set.

**Fig 6.**
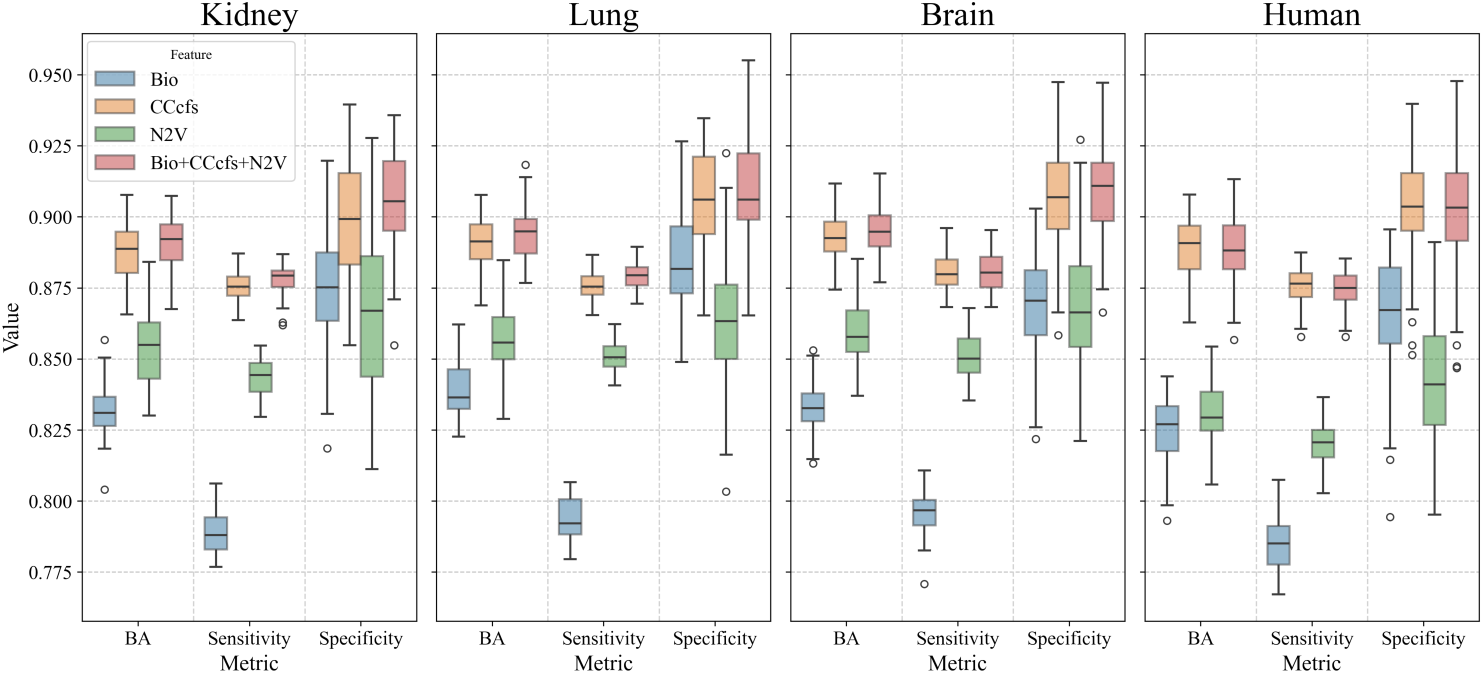
Classification performance based onHELP labelling. Box plots of the performance metrics obtained for classification “E vs NE” using different sets of features on Kidney, Lung, Brain tissues and Human.

#### Prediction performance on ucsEGs

To evaluate the ability of the model to precisely predict the csEGs and ucsEGs, we calculated the True Positive Rates (TPRs) for both (Fig 7). ucsEGs of the three contexts and their intersections are shown in Fig 7A, colored according to the prediction results. The TPR for csEGs (Fig 7B) corresponds to the Sensitivity, whose mean is reported in Table S.5, and was particularly high for all tissue cases. The TPRs for ucsEGs (Fig 7B) were instead lower due to the strong specificity of these small sets of genes. Nevertheless, achieving 80% (Kidney) and 75% (Lung) could be considered a significant outcome, suggesting promising predictive potential. In the case of the Brain, we got a TPR of 66%.

**Fig 7.**
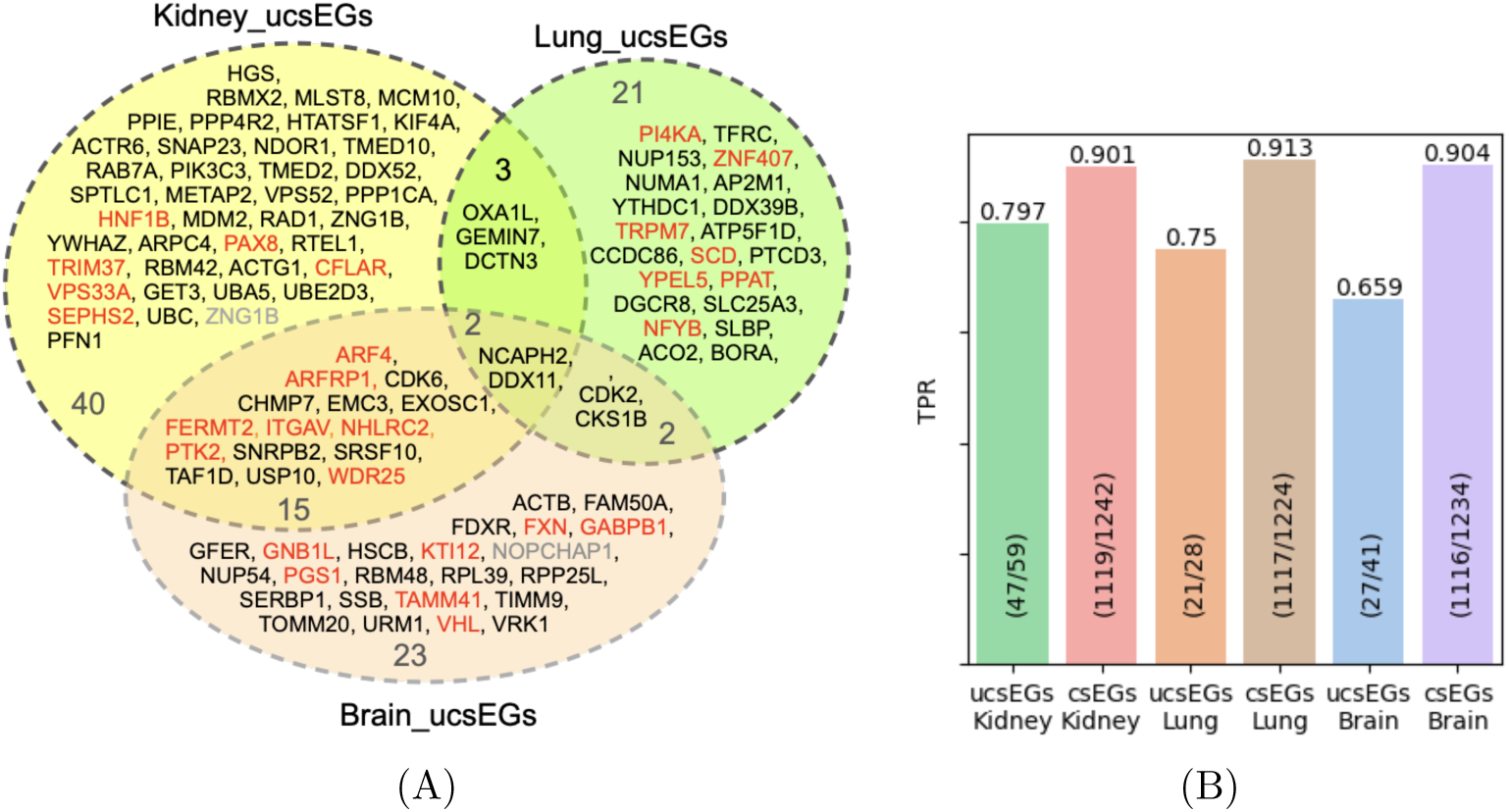
ucsEGs prediction. (A) Diagram representing the intersection of ucsEGs of Kidney, Lung and Brain tissues. The ucsEGs for each tissue are computed by subtracting the cEGs calculated over all tissues from csEGs. ucsEGs wrongly predicted are highlighted in red. Only one gene for Kidney and one for Lung (grey-colored) were not predicted as no attributes were retrieved for them. (B) TPRs of prediction of ucsEGs and csEGs computed in one cross-validation round using the optimal features set Bio+CCcfs+N2V.

#### Comparison with other predictors

Although finding similar works in the literature to compare our results was difficult due to our focus on context-specificity, we considered some of them to generally evaluate the goodness of the measures and consolidate the effectiveness of HELP. In our opinion, the approaches cannot be split by separating the model from the attributes, as these latter are a crucial part of the prediction power. For this reason, the comparisons have been conducted using the features proposed by the related works, and only the labels were uniform. In [38], the authors proposed an ML method across six model eukaryotes, called CLEARER, designed to predict cellular essential genes (CEG) and organismal essential gene (OEG), where essentiality information was collected from the OGEE [39] and DEG [40] databases. We compared the two methods (see section M.3) for the prediction of cEGs, using the OGEE+DEG annotations exploited by CLEARER’s authors as labels. We built the HELP prediction model (sveLGBM) by using as training data the Bio+CCcfs+N2V input features for the Human context, while we built the CLEARER prediction model (Random Forests) by using gene attributes and feature selection method (Lasso) as described in their work [38]. The comparison of performances (see Table S.6) proved the superiority of sveLGBM over CLEARER’s approach considering BA, ROC-AUC and Sensitivity metrics. In particular, lower Sensitivity means a low percentage of the E gene is correctly predicted. The cost to pay for a higher recall on positive samples is a lower Specificity.

DeepHE [6], which predicts Human EGs using labels from the DEG database [40] and integrating features derived from sequence data and PPI embedding, declared a ROC-AUC=0.941. EPGAT [7], a method based on Graph Attention Networks, achieved a ROC-AUC=0.915±0.0034 predicting Human EGs using labels from the OGEE database [39] to supervise the training of the GAT models and using as training data a combination of PPI and sub-localisation annotations. The three methods, i.e., sveLGBM, DeepHE, and EPGAT, were compared (see section M.4) the prediction of both cEGs and csEGs. Given the adaptation of the two literature tools to the context-specificity domain, we decided to perform the comparison using the HELP labelling in the three tissue contexts (csEGs) and Human (cEGs). The performance comparison results are reported in Table S.7 of the Supplementary Material document. They showed the superiority of sveLGBM in all the first case studies and almost all the metrics measurements. The optimal hyper-parameters setting is reported in Table S.8.

In [41], the authors proposed a model based exclusively on human PPI network embedding and reported a BA lower than ours (BA=0.783). Unfortunately, we could not investigate further the performance comparison due to the unavailability of code for this method.

### Investigating almost essentiality

One of the aims of the current work was to investigate the hypothesis of an intermediate class between E and NE. To this extent, we subdivided the class NE into two further classes, named aE (almost Essential) and sNE (strongly Not Essential), using the same procedure described in Section *Labelling*. Briefly, we applied Otsu’s thresholding to the CRISPR scores of the NE genes for each cell line and then computed the tissue-specific gene labels as the mode of the aE or sNE labels obtained for the cell lines belonging to the tissue.

We analysed the classification performance on the three binary problems “E vs sNE”, “E vs aE”, and “aE vs sNE” on Kidney tissue by using the optimal input features configuration Bio+CCcfs+N2V (Fig 8; Table S.9). As expected, discriminating E from sNE genes resulted in better performance (BA=0.915±0.007) than considering NE all the genes that are not E (BA=0.890±0.009 see Table S.5). The most challenging problem was, as expected, the separation of the aE class from the other two (Fig 8; second and third column of Table S.9), given the intermediate nature of these genes.

**Fig 8.**
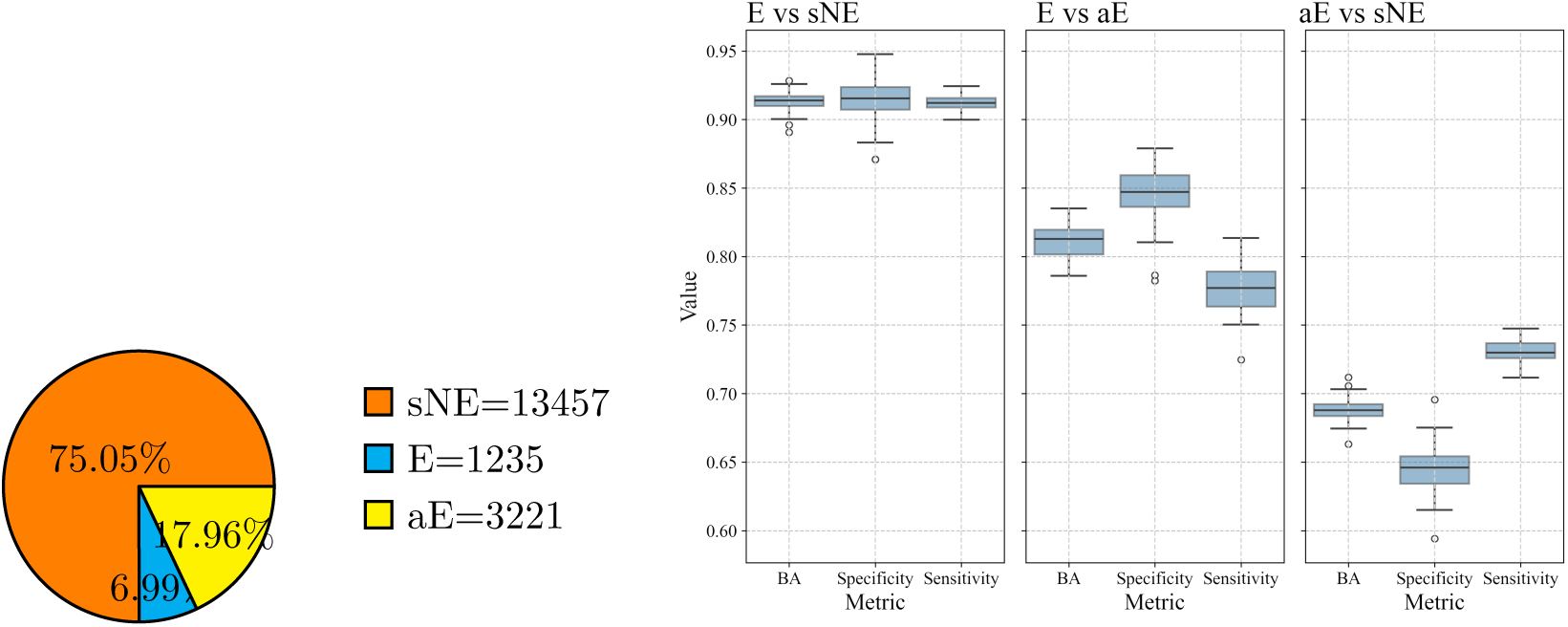
Classification performance on Kidney based on the three-class HELP labelling. (A) Three-class labelling distribution for Kidney csEGs, and (B) Box plots of the performance metrics obtained for classification of “E vs sNE”, “E vs aE”, and “aE vs sNE” using Bio+CCcfs+N2V features.

The separation of E and aE genes, although not optimal like E vs sNE, was achieved with a BA=0.813±0.012. A more pronounced reduction of the performance was observed addressing the “aE vs sNE” problem (BA=0.687±0.010).

To get valuable insights about the three classes of genes, particularly the intermediate aE, we investigated their profiles, analysing principally the attributes that contribute to their discrimination (Bio+CCcfs+N2V). These genes showed a different localisation in the PPI network (Fig 9, left side). As the layout used to represent the networks is force-directed and tends to draw nodes with greater centrality to more central positions, the resulting figures strengthen the centrality-lethality rule. The E genes (blue nodes) essentially colocalise at the centre of the network, the aE genes (yellow nodes) mostly occupy the surrounding area, and the sNE genes (orange nodes) are distributed all around, with a sort of radial pattern. Looking at the GO-CC terms enrichment (Fig 9, right side), E genes mainly enriched the nuclear body, the aE genes the mitochondrion and, to a lesser extent, the nucleus, and the sNE genes the plasma membrane and intra- and extra-cellular transport systems (e.g. granule, lysosomes, membrane rafts). Furthermore, it was interesting to notice that while E and aE genes shared some terms, even with different ranking positions, none shared any term with the sNE genes. It is also worth noting that the Bio attributes were all but one statistically different between E and sNE genes, remarking their profound difference (Fig S.5, S.6). Comparing the aE class with the other two, the situation did not change basically; 15/18 generic attributes and all the context-specific attributes were statistically different in the contrasts aE vs E and aE vs sNE, confirming the hypothesis that the aE genes were not from the same population of the other two groups. Some similarities were also present. E and aE genes, for example, showed comparable gene lengths and Driver_Genes profiles, while aE and sNE showed analogous numbers of GO-MF, GO-BP and KEGG annotations. aE and sNE genes were both extracted from the larger NE group. To demonstrate that the differences observed were not due to an implicit variability of NE genes, we extracted 100 random partitions of 3000 genes (the approximative number of aE genes) from the sNE group. We compared them to the rest of the sNE genes. Fig S.7A shows that rarely the differences were significant and, even if so, the p-value indicating the significance level was always lower than the one obtained when comparing aE. Furthermore, the same partition never shows significance for more than 5 attributes (Fig S.7B), while aE was significantly different in all but three. Moreover, dosing different percentages of aE mixed with sNE genes (to 3000 genes) and performing 10 iterations of the Wilcoxon test to compare the mean to the rest of sNE genes for some attributes, we observed a trend of the p-values, the more the percentage of aE genes in the partition, the lower the p-value (Fig S.7C). We enriched the Gene Families (gf) and GO-BP to demonstrate further that aE genes represent a distinct category. The intersection of the enriched terms for the three classes among the three tissue contexts (Fig S.8A-S.8C; S.8E-S.8G) and among the three classes for the same context (Fig S.8D, S.8H) showed a high overlapping and coherence of the characteristics of the 3 classes of genes changing the context and a very poor overlapping among the three classes, fixing the context. The gf and GO-BP terms enriched confirmed what we observed for GO-CC enrichment, assigning roles and involvement in nuclear, mitochondrial and cytoplasmic/membrane complexes and processes to E, aE and sNE genes, respectively (Supplementary file 2).

**Fig 9.**
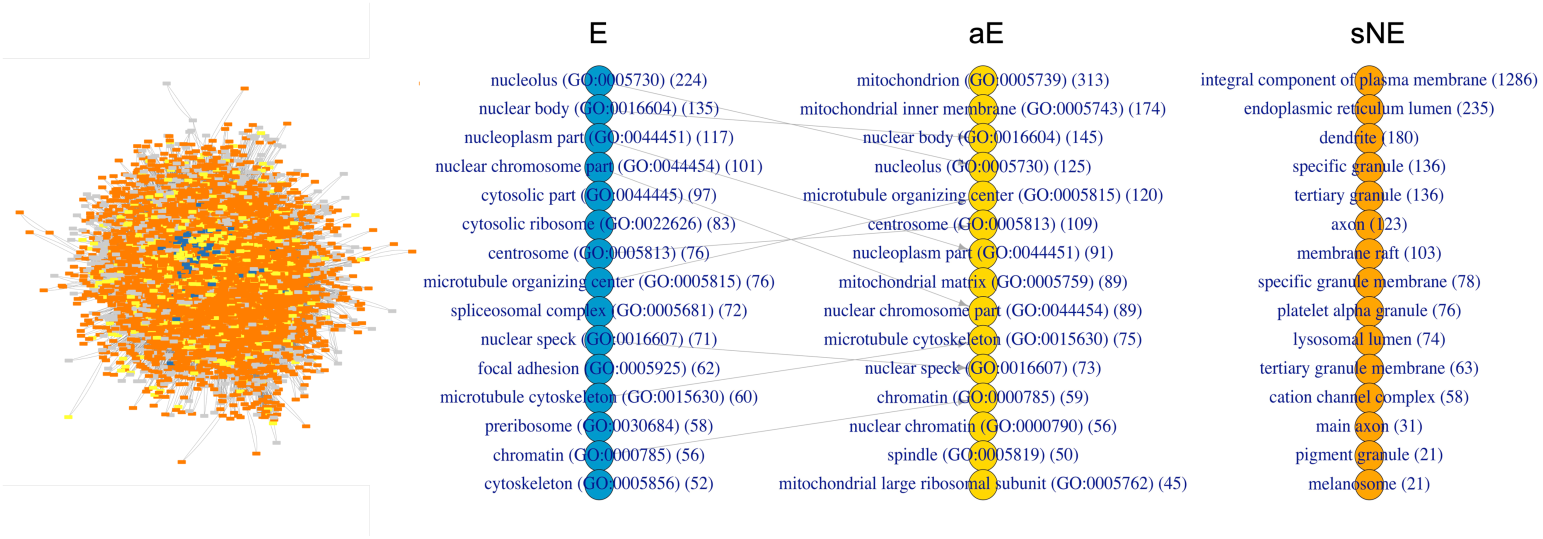
Cellular localisation of E, aE and sNE genes. The cellular sublocation of the three classes of genes is shown through the human PPI network representation (left side) and the top-ranked enriched GO-CC terms (right side) for the human context. E, aE and sNE genes are highlighted in blue, yellow and orange, respectively, in both graphics. In favour of visualisation, only the largest connected component of the network is shown. Grey nodes are genes in the PPI not labelled by HELP since they are not included in the experimental data. The network was plotted through Cytoscape v3.10.1 [42]. The rank-plots (right-side) show the top 15 significant terms ordered by gene count. The number was chosen because of fifteen significant terms enriched by sNE genes. The number of genes enriching each term is in brackets. The connection lines indicate the terms shared by the groups. The enrichment analysis was performed using the enrichR package v3.2 [43]. The list of genes with annotation of the group is reported in the Supplementary file 1.

## Discussion

This work was conceived to address two of the main issues concerning csEGs identification: labelling and prediction. The first regards the assignment of E/NE labels using the data furnished by gene deletion experiments. The need to create personalised contexts and the incessant updates of the experiments require a flexible labelling method to apply to the knockout experimental scores. The second issue regards predicting those labels by training ML methods with gene attributes from multi-source data.

We presented the HELP framework and demonstrated its effectiveness on organism-wide context, three human tissues and two subtypes of lung cancer. We applied an unsupervised thresholding method to derive both cEGS and csEGs from CRISPR scores. The obtained labels were validated by comparing the resulting cEGS with the CoRe’s methods, ADaM and FiPer, and with four reference sets of cEGs. The overlap with these sets was particularly high, especially in the case of the most recent ones (Fig 3). The results make HELP the best compromise between the overlap with reference sets and false positive rates, with the advantage over FiPer being more stringent and over ADaM being unsupervised. The identification of cEGs allows to better capture the context specificity by subtracting from the csEGs the genes essential in most contexts. This subtraction step gives rise to what we call ucsEGs. The stringency is crucial to guarantee higher confidence when approaching the context-specificity. The discrimination of the “essentiality” classes was performed by training a prediction model through knowledge from multi-omics data and embedding a context-specific PPI. As evident, the classification faces a strong unbalanced issue, where the majority class can be 13 times bigger than the minority. This strongly affects classification performance, resulting in high specificity and low sensitivity. To this extent, we propose a new meta-learning model, namely Splitting Voting Ensemble (SVE), which is based on a soft-voting ensemble of classifiers, each one trained on an equal portion of the majority class set based plus the rest of the dataset where the number of ensemble members was given by partitioning the majority class set based on the data distribution. This approach reduced the unbalancing, still using the whole dataset, in contrast to the subsampling, without duplicating data, as done by the oversampling, and without introducing synthetic data, as in the augmentation. The best-performing configuration was given by combining two sets of features, Bio+CCcfs, with the addition of the embedding features N2V for better addressing the context-specificity. The Bio set contains a collection of generic and context-specific functional and structural characteristics; the CCcfs set contains information on cellular localisation and involvement in molecular complexes; the N2V set contains topological information from the PPI, which can be context-specific. The feature importance analysis highlighted that the Bio attributes concerning interactions ("BIOGRID"), context-specific gene expression ("OncoDB expression", "HPA", "GTEX"), and some physical ("Gene length", "GC content") and conservative ("Orthologs count") properties are the most discriminative between E and NE (Fig S.9). Among the CCcfs annotations, the most important are the involvements in spliceosomal and polymerase complexes, properties of EGs, as suggested by the enrichment of gene families and biological processes (Supplementary file 2). The localization in cellular compartments, nucleus, mitochondrion, and membrane also showed high importance when considering the CCcfs attributes. The model could correctly predict around 90% of csEGs and between 66%-80% of ucsEGs. This discrepancy was likely due to the inherent challenge in modelling the context-specificity of EGs, stemming from their small number compared to the csEGs and the limited availability of context-specific information characterising them accurately.

Several recent papers discussing the characteristics of EGs [11, 44] reveal that essentiality is not a binary fixed but a flexible status depending on the genetic and environmental contexts. The characteristics predictive of essentiality are quantitative traits, as are the essentiality scores. In this scenario, it is reasonable to think that the dividing line between essentiality and not is uncertain and that a gene essential in some contexts likely keeps involvement in essential functions in others. In our previous works, using a different labelling strategy based on a knowledge-driven subdivision of CRISPR scores (CS), we identified the best configuration of E and NE genes by ML trials. We achieved the best performance training a model on biological and embedding attributes (CS0 (E) vs CS6-9 (NE): BA=0.84 [45, 46]). The investigation of an intermediate group led us to consider possibly overcoming the dichotomic view of essentiality. The experimental evidence we got, along with the above considerations, suggest the existence of shades of “essentiality”, that, in the most simplistic view, can be represented by a third class of “almost Essentials” (aE). Our prediction results demonstrated that the attributes we collected for classifying the genes are a quantitative reflection of the essentiality, given the performance of the classification and the improvement we got when trying to classify the most extreme groups of genes, E and sNE. This means that the more distant the knockout scores, the more our attributes separate the genes. Most features in the Bio set have values statistically different between aE and the other two groups, E and sNE (Figs S.5, S.6). Another piece of evidence for considering aE as a distinct class is the analysis of these genes’ functional and physical involvement. Different gene families (gf), biological processes (GO-BP) and cellular localisation (GO-CC) are shown by the three classes of genes, with few overlapping between E and aE (giving reason to the name "almost Essential") and no overlapping between aE and sNE, although they derive from the same larger group NE. Changing the context, the characteristics of the three classes are maintained (Fig S.8). E genes mostly enriched the nucleus area (GO-CC), the ribosomal, proteasome, polymerase families (gf), and the replication machinery (GO-BP), while aE genes mostly enriched the mitochondrion (GO-CC), its proteins (gf) and processes (GO-BP) (Supplementary File 2). This result highlights the promising role of the aE as candidate biomarkers, as the mitochondria are considered not only powerhouses but also dynamic regulators of life, death, proliferation, motion and stemness of cancer cells [47].

Moreover, this perfectly aligns with the role of mitochondria to fill the functional gap between the nucleus and cytoplasmic organelles and the interconnection between their components [48]. sNE genes, instead, enriched the plasma membrane and cellular transportation systems (GO-CC), cell surface molecules (gf) and cell adhesion and communication systems (GO-BP) (Supplementary File 2). The different localisation was also evident by visualising the three groups in the PPI network, which seems to reproduce the picture of a human cell, with the E genes localised in the network’s core, aE in the surrounding area and sNE widely spread at the borders (Fig 9). Our recent work demonstrated that integrating the PPI with a metabolic network to add a functional centrality to the physical one, the contribution was totally in charge of the PPI [46]. This can likely be explained by the fact that while the metabolic machinery comprises several alternative paths to achieve a specific objective, the lack of a component involved in many physical complexes and interactions is hard to tolerate.

A fair and complete examination of the proposed model requires a discussion of its limitations. For HELP, most of them regard the context-specific approach since it is a more unexplored topic. First of all, the lack of experimental validation for csEGs automatically determines the absence of ground truth to validate the model besides the labels produced by the workflow and the difficulty of comparing our method to others without adapting the provided codes and data. The usability of HELP for context-specific investigations is strictly related to data availability. As an example, the main limitation of the labelling process is the availability of sufficient gene deletion score data. The lower the number of cell lines, the less significant the mode computation over a small sample size of partial labels. Public context-specific attributes are generally limited, but HELP’s flexibility has been thought to manage custom data. Some sources used in this paper that contributed to the results presented could not contain data for some contexts and should be replaced by different sources. For example, HPA and GTEX data must be considered as gene expression annotations that are not necessarily linked to a specific source. The context-specific PPIs used to extract gene features by deep learning in embedding vectors are built considering the gene expression in the specific tissue based on experimental, orthology or prediction evidence and not on the real physical interactions experimentally evaluated in the specific context. This implicitly involves limits regarding the reliability of the physical connections. Furthermore, retrieving a PPI for each specific context of interest is impossible, but the user can choose to use the generic human PPI or the one that can be considered closer to the case study. Currently, the prediction method does not use feature selection techniques to reduce the input data size. The feature selection adopted can be defined as knowledge-driven, as the attributes potentially predictive of the essentiality have been collected by examining the literature on the topic. However, the only high-sized feature set is CCcfs. The experiments we conducted on feature importance (see Section M.5 in Supplementary Methods document) show that these features taken all together provide the larger contribution to the classification, and this is reinforced by the finding that the reduction of this feature set does not determine any performance improvements but rather a degradation of 1-2 % in terms of average BA (over ten iterations of the experiments). Of course, if we consider additional features not included in the current study, feature selection might also represent a valid approach for performance improvements. The prediction model sveLGBM, although outperforming, is time costly: we pay a timing overhead in executing an ensemble of LGBM classifiers, each one of them being by itself an ensemble of decision trees. In our opinion, this cost is reasonable for the gain obtained in performance.

Despite the cited limitations, we strongly believe that the promising insights extracted open the way to future work to investigate the essentiality shades and apply HELP to precision medicine purposes. In this regard, we here demonstrated its application to a disease case study, identifying the EGs specific to two different lung cancer subtypes, NSCLC and NET, that can represent ideal candidates for a precise recognition and targeting of cancer cells. The stringency of HELP in considering NSCLC and NET ucsEGs is particularly advantageous as a small number of candidates are manageable and can benefit disease studies. Furthermore, the genes identified showed some interesting traits: NET ucsEGs are involved in cellular respiration and energy production (Fig S.3), mechanisms particularly associated with NETs [36], while the eight NSCLC ucsEGs are all significantly differentially expressed comparing the two NSCLC subtypes (LUAD and LUSC) with the normal samples (Fig S.4). The novelty of HELP is also represented by the attributes used for EGs prediction. To the best of our knowledge, and as demonstrated by the comparison we performed, most of the tools aimed at predicting EGs use few and recurrent attributes, such as PPI, gene expression, orthologs count, and sequence information. Although, in literature, some characteristics that we used as attributes have been associated with EGs, they have never been used for their classification (e.g. BIOGRID, UP_tissue, Driver genes, Gene-Disease association, TFBs). In this scenario, the attributes "BIOGRID", "REACTOME", and "UCSC_TFBS", which annotate the functional interactions of genes, the involvement in pathways and the transcription factors binding site predictions, respectively, are among those showing the highest scores of importance (Figure S.9). Investigating the diversified attributes that predict these genes has suggested new insights about their cellular localization and functions. Last but not least, we would like to highlight that the two strategies of HELP, labelling and prediction, which are distinct even if connectable, serve a double scope: the labelling, also considered an identification method, can help to process and rationalise the experimental knock-out results and are therefore strictly dependent on the experiments, while the ML and DL approaches can help in establishing the essentiality traits to recognise essential genes in different contexts, not only supporting but potentially substituting massive wet lab genome-wide experiments, which are not void of technical biases [49]. The learning models designed not only represent a valuable tool for EGs prediction but may also be considered as an additional strategy for the validation of labelling: a high performance in predicting HELP-based labels suggests that those labels are well-fitted by learning models trained on gene features known to represent facets of the concept of essentiality.

## Acknowledgments

This work was carried out also within the activities of the authors as members of the INdAM Research group GNCS and the ICAR-CNR INdAM Research Unit. Authors from ICAR-CNR would like to thank R. Mattiello, G. Trerotola and S. Sada for their technical support.

## Supporting information

### Data/Software Availability

Gene Effect file (DepMap v.23Q4) and Model are third-party data used in this work that can be retrieved from https://doi.org/10.25452/figshare.plus.24667905.v2. All other data and software produced in this work are released under the GNU licence and are available in three repositories:

- https://zenodo.org/doi/10.5281/zenodo.10964743 is the Zenodo DOI for downloading the GitHub repository (https://github.com/giordamaug/HELP) storing all data produced in the current study, i.e. gene labelling files, context-specific EGs lists, EG prediction results and performance measurements in each tissue/disease context. The repository includes a directory of notebooks to carry on all processing and experiments discussed in the manuscript: the extraction of PPI embeddings (embedding.ipynb), the identification of csEGs (labelling.ipynb) and ucsEGs (csegs.ipynb), the prediction of csEGs (prediction.ipynb), the importance analysis of gene features used to build prediction models (feature_importance.ipynb), the hyper-parameters optimization of the prediction model (optuna.ipynb) and the comparison with alternative prediction methods (compare_models.ipynb).
- https://doi.org/10.5281/zenodo.12597679 is a Zenodo repository containing only input data used for the classification of essential genes in tissue-specific contexts. These data are huge and stored in a separate repository since we wanted to provide users with a lighter downloadable repository for only the HELP software and tools. In addition, the sizes of these files overcome the limits of the GitHub archiving facility.
- https://doi.org/10.5281/zenodo.12598244 is the Zenodo DOI for downloading the GitHub repository (https://github.com/giordamaug/SVElearn) where you find source code and examples of usage of the Splitting Voting Ensemble approach (SVE). We designed and developed this machine learning method to provide high classification performance in cases of highly unbalanced datasets. Therefore, its implementation and application go beyond its specific use in the domain of essential gene prediction, which is the focus of this work.

## Supplementary Material

**Fig S.1. ucsEGs PPI enrichment**. PPI networks built through STRING [50] using the ucsEGs computed for Kidney (A), Lung (B) and Brain (C). The nodes are coloured according to the enriched terms shown in the associated tables. The significant (False Discovery Rate, FDR *<* 0.05) non-redundant terms were ranked by the number of enriching genes (Count in the network: no. of enriching genes/no. of genes annotated for the term). The edges were built with all the STRING information except “Text mining".

**Fig S.2. Disease-specific ucsEGs.** Diagram representing disease-specific (Non-Small-Cell Lung Cancer NSCLC and Lung Neuroendocrine Tumour NET) and lung ucsEGs intersections by ADaM, FiPer, and HELP labelling. Each row represents the set of ucsEGs for each labelling. The last row reports the number of genes resulting from the intersections. The last column on the right indicates the number of ucsEGs for each set, with the dark grey shadow representing the corresponding histogram.

**Fig S.3. Reactome pathway enrichment of lung NET-specific EGs.** The significantly enriched pathways are shown on the y axis; the color bar indicates the significance in terms of False Discovery Rate (FDR)-adjusted p-value, while the dot size indicates the number of genes in the input set found in the pathway. On the x axis the Fold Enrichment, namely the percentage of genes in the input list annotated in a pathway divided by the corresponding percentage in the background human genes.

**Fig S.4. Differential expression of NSCLC ucsEGs.** The boxplots show the expression levels of the eight NSCLC-specific EGs in the two NSCLC subtypes, LUAD and LUSC, and normal samples, as collected in OncoDB. The significance of the average difference between the two populations was evaluated with a Student’s t-test using the OncoDB platform tool for the differential expression analysis. The legends indicate the colours associated with the groups and the number of samples in brackets.

**Fig S.5. Boxplots of the generic Human Bio attribute values for the E, aE, and sNE classes.** The stars on the top indicate the significance of the Wilcoxon test for each pair of comparisons (**** *≤* 0.0001, *** *≤* 0.001, ** *≤* 0.01, * *≤* 0.05, ns = not significant). In favour of visualisation, the values have been signed-square-root transformed.

**Fig S.6. Boxplots of the context-specific Bio attribute values of the three tissues investigated for the E, aE, and sNE classes.** The stars on the top indicate the significance of the Wilcoxon test for each pair of comparisons (**** *≤* 0.0001, *** *≤* 0.001, ** *≤* 0.01, * *≤* 0.05, ns = not significant). The Driver genes attributes were not shown as having small ranges of values and poor statistics. In favour of visualisation, the values have been signed-square-root transformed.

**Fig S.7. Random extraction of the intermediate class.**. A) For each generic attribute (taken as an example from the Kidney dataset) and cs attributes from the three tissues, 100 random partitions of 3000 genes from the sNE groups have been extracted and compared to the rest of the sNE genes. For each tissue, the 100 partitions were fixed. Wilcoxon test was performed to evaluate the statistical significance (p-value) and verify whether the groups come from the same population for each pair of comparisons (**** *≤* 0.0001, *** *≤* 0.001, ** *≤* 0.01, * *≤* 0.05, ns = not significant). The table indicates the number of partitions for each attribute and for each significance level indicated in the column header. The level of significance given by comparing aE vs sNE, and indicated in Figures S.5;S.6, was also shown by the orange text "aE". B) The histogram shows the number of attributes (x-axis) for which the partitions are simultaneously significant. The count of partitions (y-axis) for each frequency is also shown on the bars. C) The line plot shows the mean of −log10(p-value) and the standard deviation from Wilcoxon tests between different percentages of aE mixed with sNE genes (to 3000 genes) obtained with 10 iterations and the rest of sNE genes for some attributes indicated in the legend.

**Fig S.8. Intersection of Gene Families and Biological Processes enrichment among E, aE and sNE genes.** The Venn diagrams show the intersection of Gene Families (gf) and Gene-Ontology Biological Processes (BP) enriched by E, aE or sNE genes among the three tissue contexts under study (A-C; E-G), as well as the intersection of Gene Families (gf) and Gene-Ontology Biological Processes (BP) enriched by genes of the three classes in one context (here Kidney tissue as example) (D;H). The number of genes composing each set is shown in brackets.

**Fig S.9. Feature importance analysis**. Bio+CCcfs attributes importance calculated by training a sveLGBM model on the entire dataset. The plot cuts-off feature with importance lower than 0.25 %.

**Table S.1. Collected genomic, transcriptomic, epigenetic, functional and evolutionary features of genes.** (cs) indicates the context-specific attributes.

**Table S.2. Comparison of classifiers on prediction in “E vs NE” problem in the Kidney case study.** Ranking of methods is based on the Balanced Accuracy metric. All methods with “sve” prefix are our meta-learning model proposal with a different base classifier as member of the ensemble. All other methods are provided by the PyCaret library. All models where trained with Bio+CCcfs+N2V attributes of genes. CPU times are measured on Apple M2 with 16GB RAM.

**Table S.3. sveLGBM tuning of parameters with Optuna library**. Optimiziation was carried out on “E vs NE” classification problem with a stratified 5-fold cross-validation with Bio+CCcfs+N2V features by maximising BA metric.

**Table S.4. Classification performance metrics adopted in the experiments.** They are defined in terms of the number of true positives (TP), true negatives (TN), false positives (FP), and false negatives (FN), where the first class in each binary task (e.g. class E in the “E vs NE” classification task) is assumed as the positive class.

**Table S.5. “E vs NE” classification performance based on HELP labelling.** (A) Kidney, (B) Lung, (C) Brain tissues, and (D) Human. Averages and errors of metrics are obtained on fifty measurements related to ten times iterated 5-fold cross-validation. The averaged Confusion Matrix (CM) is also shown.

**Table S.6. Comparison of sveLGBM and CLEARER on OGEE+DEG labelling for the prediction of cEGs.** *Hs Features* refer to the features collected for Homo Sapiens EGs prediction presented in the work [38]. sveLGBM hyperparameters: n_voters=16, learning_rate=0.1, n_estimators=200, boosting_type=’gbdt’.

CLEARER hyperparameter: RF n_estimators=500 as in [38].

**Table S.7. Comparison of sveLGBM, DeepHE and EPGAT predictions on HELP labelling for Kidney-, Lung-, Brain-specific EGs, and cEGs (Human).** EPGAT running with PPI input and sublocalisation attributes. EPGAT hyper-parameters are optimised by using the provided tuning function. DeepHE running with DNA sequencing extracted features plus node2vec embedding 120-sized features extracted from the PPI. HELP running with Bio+CCcfs + N2V embedding 120-sized features extracted from the PPI.

**Table S.8. Optimal hyper-parameters of sveLGBM, DeepHE and EPGAT methods used in comparison of Table S.7.**

**Table S.9 “E vs sNE”, “E vs aE” and “aE vs sNE” classification performance based on HELP labelling.** The case study is Kidney tissue using Bio+CCcfs+N2V features. Averages and errors of metrics are obtained on fifty measurements related to ten times iterated 5-fold cross-validation. The averaged Confusion Matrix (CM) is also shown.

## Supplementary Methods

M.1 Base estimator choice for SVE. Description of the experiments aimed at tuning and comparing the performance of several classifiers on the binary classification problem E/NE.
M.2 sveLGBM classifier tuning. Hyper-parameters optimisation of sveLGBM.
M.3 Comparison with CLEARER. Comparison of sveLGBM with CLEARER for the prediction of cEGs.
M.4 Comparison with DeepHE and EPGAT. Comparison of sveLGBM with DeepHE and EPGAT methods for the prediction of cEGs and csEGs.
M.5 Feature importance analysis. Analysis of features’ importance in E/NE genes classification.

## Supplementary files

**Supplementary file 1** Excel file containing the gene names and the associated HELP labelling E/aE/sNE used to assign colours in Figure 9.

**Supplementary file 2** Excel file containing the results of Gene Families (gf) and Biological Processes (GO-BP) enrichment. The first was obtained by downloading gene families annotation from https://www.genenames.org/download/statistics-and-files/, applying a hypergeometric test (R version 4.1.2) using all the genes in the DepMap matrix as background. The GO-BP enrichment, instead, was performed by using DAVID Bioinformatics tool (https://david.ncifcrf.gov/tools.jsp). Each sheet is named according to the content “tissue_enrichment_class". The columns content is detailed in each sheet.

## References

1. Juhas M, Eberl L, Glass JI. Essence of life: essential genes of minimal genomes. Trends in cell biology. 2011;21(10):562–568.

2. Gurumayum S, et al. OGEE v3: Online GEne Essentiality database with increased coverage of organisms and human cell lines. Nucleic Acids Res. 2021;49(D1):D998–D1003.

3. Ferreira P, Choupina AB. CRISPR/Cas9 a simple, inexpensive and effective technique for gene editing. Mol Biol Rep. 2022;49(7):7079–7086.

4. Aromolaran O, Aromolaran D, Isewon I, et al. Machine learning approach to gene essentiality prediction: a review. Brief Bioinform. 2021;22(5):bbab128.

5. Giordano M, Falbo E, Maddalena L, Piccirillo M, Granata I. Untangling the Context-Specificity of Essential Genes by Means of Machine Learning: A Constructive Experience. Biomolecules. 2024;14(1).

6. Zhang X, Xiao W, Xiao W. DeepHE: Accurately predicting human essential genes based on deep learning. PLoS Comput Biol. 2020;16(9):e1008229.

7. Schapke J, et al. EPGAT: Gene Essentiality Prediction With Graph Attention Networks. IEEE/ACM Trans Comput Biol Bioinform. 2022;19(3):1615–1626.

8. Le NQK, Do DT, et al. A computational framework based on ensemble deep neural networks for essential genes identification. Int J Mol Sci. 2020;21(23):9070.

9. Ashtiani M, Salehzadeh-Yazdi A, Razaghi-Moghadam Z, et al. A systematic survey of centrality measures for protein-protein interaction networks. BMC Syst Biol. 2018;12(1):1–17.

10. Larrimore KE, Rancati G. The conditional nature of gene essentiality. Curr Opin Genet Dev. 2019;58:55–61.

11. Rancati G, Moffat J, Typas A, Pavelka N. Emerging and evolving concepts in gene essentiality. Nat Rev Genet. 2018;19(1):34–49.

12. Zhang W, Quevedo J, Fries GR. Essential genes from genome-wide screenings as a resource for neuropsychiatric disorders gene discovery. Translational Psychiatry. 2021;11(1):317.

13. Dickerson JE, Zhu A, Robertson DL, Hentges KE. Defining the role of essential genes in human disease. PloS one. 2011;6(11):e27368.

14. Vinceti A, Karakoc E, et al. CoRe: a robustly benchmarked R package for identifying core-fitness genes in genome-wide pooled CRISPR-Cas9 screens. BMC Genom. 2021;22(828).

15. Sharma S, Dincer C, Weidemüller P, et al. CEN-tools: an integrative platform to identify the contexts of essential genes. Mol Syst Biol. 2020;16(10):e9698.

16. Behan FM, Iorio F, Picco G, et al. Prioritization of cancer therapeutic targets using CRISPR-Cas9 screens. Nature. 2019;568(7753):511–516.

17. Hart T, Tong AHY, Chan K, et al. Evaluation and Design of Genome-Wide CRISPR/SpCas9 Knockout Screens. G3 Genes|Genomes|Genetics. 2017;7(8):2719–2727.

18. Otsu N. A Threshold Selection Method from Gray-Level Histograms. IEEE Trans Syst Man Cybern. 1979;9(1):62–66.

19. Szklarczyk D, Kirsch R, Koutrouli M, Nastou K, Mehryary F, Hachilif R, et al. The STRING database in 2023: protein–protein association networks and functional enrichment analyses for any sequenced genome of interest. Nucleic acids research. 2023;51(D1):D638–D646.

20. Kotlyar M, Pastrello C, Ahmed Z, et al. IID 2021: towards context-specific protein interaction analyses by increased coverage, enhanced annotation and enrichment analysis. Nucleic Acids Res. 2021;50(D1):D640–D647.

21. Grover A, Leskovec J. Node2vec: Scalable Feature Learning for Networks. In: KDD ‘16; 2016. p. 855–864.

22. Mikolov T, Sutskever I, et al. Distributed Representations of Words and Phrases and Their Compositionality. In: Proc. NIPS’13 - Vol.2; 2013. p. 3111–3119.

23. Ardlie KG, et al. The Genotype-Tissue Expression (GTEx) pilot analysis: Multitissue gene regulation in humans. Science. 2015;348(6235):648–660.

24. Sherman BT, Hao M, Qiu J, Jiao X, Baseler MW, Lane HC, et al. DAVID: a web server for functional enrichment analysis and functional annotation of gene lists (2021 update). Nucleic Acids Res. 2022;50(W1):W216–W221.

25. Tang G, Cho M, Wang X. OncoDB: an interactive online database for analysis of gene expression and viral infection in cancer. Nucleic Acids Res. 2022;50(D1):D1334–D1339.

26. Uhlén M, Fagerberg L, Hallström BM, et al. Tissue-based map of the human proteome. Science. 2015;347(6220).

27. Binder JX, Pletscher-Frankild S, Tsafou K, et al. COMPARTMENTS: unification and visualization of protein subcellular localization evidence. Database. 2014;2014.

28. Oughtred R, Stark C, Breitkreutz BJ, et al. The BioGRID interaction database: 2019 update. Nucleic Acids Res. 2019;47(D1):D529–D541.

29. Sayers EW, Bolton EE, Brister JR, Canese K, Chan J, Comeau DC, et al. Database resources of the National Center for Biotechnology Information in 2023. Nucleic Acids Res. 2023;51(D1):D29–D38.

30. Liu SH, Shen PC, Chen CY, et al. DriverDBv3: a multi-omics database for cancer driver gene research. Nucleic Acids Res. 2020;48(D1):D863–D870.

31. Piñero J, Ramírez-Anguita JM, Saüch-Pitarch J, Ronzano F, Centeno E, Sanz F, et al. The DisGeNET knowledge platform for disease genomics: 2019 update. Nucleic Acids Res. 2020;48(D1):D845–D855.

32. Giordano M, Granata I. ICARlearn: ensembling methods for efficient prediction of unbalanced data;. Available from: https://github.com/giordamaug/ICARlearn.

33. Breiman L. Random Forests. Machine Learning. 2001;45(1):5–32. doi:10.1023/A:1010933404324.

34. Schapire RE. In: Schölkopf B, Luo Z, Vovk V, editors. Explaining AdaBoost. Berlin, Heidelberg: Springer Berlin Heidelberg; 2013. p. 37–52. Available from: 10.1007/978-3-642-41136-6_5.

35. Ke G, Meng Q, Finley T, et al. LightGBM: A Highly Efficient Gradient Boosting Decision Tree. In: Proc. NIPS’17; 2017. p. 3149–3157.

36. Soldevilla B, López-López A, Lens-Pardo A, Carretero-Puche C, Lopez-Gonzalvez A, La Salvia A, et al. Comprehensive plasma metabolomic profile of patients with advanced neuroendocrine tumors (NETs). Diagnostic and biological relevance. Cancers. 2021;13(11):2634.

37. Thölke P, Mantilla-Ramos YJ, Abdelhedi H, Maschke C, Dehgan A, Harel Y, et al. Class imbalance should not throw you off balance: Choosing the right classifiers and performance metrics for brain decoding with imbalanced data. NeuroImage. 2023;277:120253.

38. Beder T, Aromolaran O, Dönitz J, et al. Identifying essential genes across eukaryotes by machine learning. NAR Genom Bioinform. 2021;3(4):lqab110.

39. Chen WH, Lu G, Chen X, Zhao XM, Bork P. OGEE v2: an update of the online gene essentiality database with special focus on differentially essential genes in human cancer cell lines. Nucleic Acids Res. 2016;45(D1):D940–D944. doi:10.1093/nar/gkw1013.

40. Luo H, Lin Y, Liu T, Lai FL, Zhang CT, Gao F, et al. DEG 15, an update of the Database of Essential Genes that includes built-in analysis tools. Nucleic Acids Res. 2021;49(D1):D677–D686.

41. Dai W, Chang Q, Peng W, Zhong J, Li Y. Network embedding the protein–protein interaction network for human essential genes identification. Genes. 2020;11(2):153.

42. Shannon P, Markiel A, Ozier O, et al. Cytoscape: a software environment for integrated models of biomolecular interaction networks. Genome Res. 2003;13(11):2498–2504.

43. Kuleshov MV, Jones MR, Rouillard AD, et al. Enrichr: a comprehensive gene set enrichment analysis web server 2016 update. Nucleic Acids Res. 2016;44(W1):W90–W97.

44. Chen H, Zhang Z, et al. New insights on human essential genes based on integrated analysis and the construction of the HEGIAP web-based platform. Brief Bioinform. 2020;21(4):1397–1410.

45. Manzo M, Giordano M, et al. Novel Data Science Methodologies for Essential Genes Identification Based on Network Analysis. In: Data Science in Applications. Springer; 2023. p. 117–145.

46. Granata I, Giordano M, et al. Network-Based Computational Modeling to Unravel Gene Essentiality. In: Mondaini RP, editor. Trends in Biomathematics: Modeling Epidemiological, Neuronal, and Social Dynamics. Springer Nature Switzerland; 2023. p. 29–56.

47. Grasso D, Zampieri LX, Capelôa T, et al. Mitochondria in cancer. Cell stress. 2020;4(6):114.

48. Gasparotto M, et al. Nuclear and Cytoplasmatic Players in Mitochondria-Related CNS Disorders: Chromatin Modifications and Subcellular Trafficking. Biomolecules. 2022;12(5):625.

49. Dede M, Kim E, Hart T. Biases and blind-spots in genome-wide CRISPR knockout screens. bioRxiv. 2020; p. 2020–01.

50. Szklarczyk D, Gable AL, Lyon D, Junge A, Wyder S, Huerta-Cepas J, et al. STRING v11: protein–protein association networks with increased coverage, supporting functional discovery in genome-wide experimental datasets. Nucleic acids research. 2019;47(D1):D607–D613.

51. Durinck S, Spellman PT, et al. Mapping identifiers for the integration of genomic datasets with the R/Bioconductor package biomaRt. Nature Protocols. 2009;4:1184–1191.

52. Ali M. PyCaret: An open source, low-code machine learning library in Python; 2020. Available from: https://www.pycaret.org.

53. Akiba T, Sano S, Yanase T, Ohta T, Koyama M. Optuna: A Next-generation Hyperparameter Optimization Framework. In: Proceedings of the 25th ACM SIGKDD International Conference on Knowledge Discovery and Data Mining; 2019.

54. Chawla NV, Bowyer KW, Hall LO, Kegelmeyer WP. SMOTE: synthetic minority over-sampling technique. J Artif Int Res. 2002;16(1):321–357.

