## Supplementary Material for "HELP: A computational framework for labelling and predicting human common and context-specific essential genes"

#### Supplementary Figures

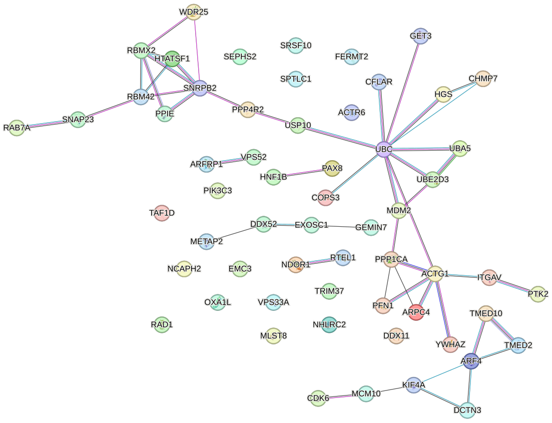


| Category | Term ID | Term description | Count in network | FDR |
|---|---|---|---|---|
| GO Component | GO:0043231 | Intracellular membrane-bounded organelle | 55/12149 | 5.73E-05 |
| GO Process | GO:0008152 | Metabolic process | 40/7988 | 0.0151 |
| GO Process | GO:0071840 | Cellular component organization or biogenesis | 38/5639 | 9.22E-05 |
| GO Process | GO:0008104 | Protein localization | 19/1943 | 0.0045 |
| GO Process | GO:0015031 | Protein transport | 15/1180 | 0.0035 |
| UniProt Keywords | KW-0597 | Phosphoprotein | 41/8122 | 0.0026 |
| Reactome | HSA-162582 | Signal Transduction | 18/2540 | 0.0233 |

(A)

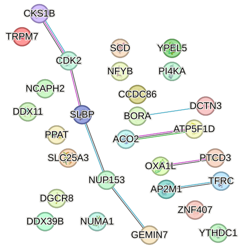


| Category | Term ID | Term description | Count in network | FDR |
|---|---|---|---|---|
| GO Component | GO:0070013 | Intracellular organelle lumen | 20/5660 | 0.0075 |
| UniProt Keywords | KW-0597 | Phosphoprotein | 22/8122 | 0.0219 |
| Reactome | HSA-69278 | Cell Cycle, Mitotic | 7/526 | 0.0167 |

(B)

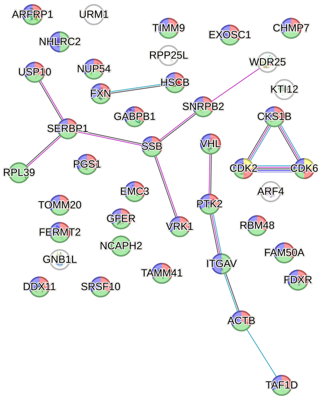


| Category | Term ID | Term description | Count in network | FDR |
|---|---|---|---|---|
| GO Component | GO:0043229 | Intracellular organelle | 35/9609 | 0.0028 |
| GO Component | GO:0043227 | Membrane-bounded organelle | 33/9083 | 0.0041 |
| GO Component | GO:0043231 | Intracellular membrane-bounded organelle | 31/8162 | 0.0041 |
| GO Component | GO:0097128 | Cyclin D1-CDK4 complex | 2/4 | 0.0206 |

(C)

**Fig S.1. ucsEGs PPI enrichment**. PPI networks built through STRING [50] using the ucsEGs computed for Kidney (A), Lung (B) and Brain (C). The nodes are coloured according to the enriched terms shown in the associated tables. The significant (False Discovery Rate, FDR < 0.05) non-redundant terms were ranked by the number of enriching genes (Count in the network: no. of enriching genes/no. of genes annotated for the term). The edges were built with all the STRING information except “Text mining".

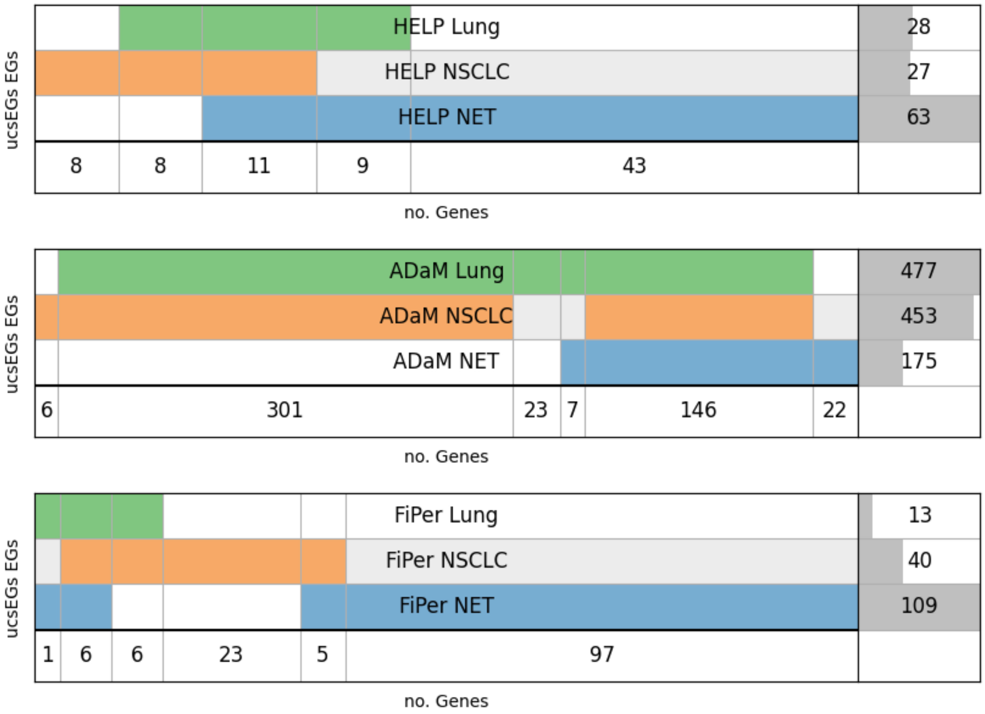


**Fig S.2. Disease-specific ucsEGs.** Diagram representing disease-specific (Non-Small-Cell Lung Cancer NSCLC and Lung Neuroendocrine Tumour NET) and lung ucsEGs intersections by ADaM, FiPer, and HELP labelling. Each row represents the set of ucsEGs for each labelling. The last row reports the number of genes resulting from the intersections. The last column on the right indicates the number of ucsEGs for each set, with the dark grey shadow representing the corresponding histogram.

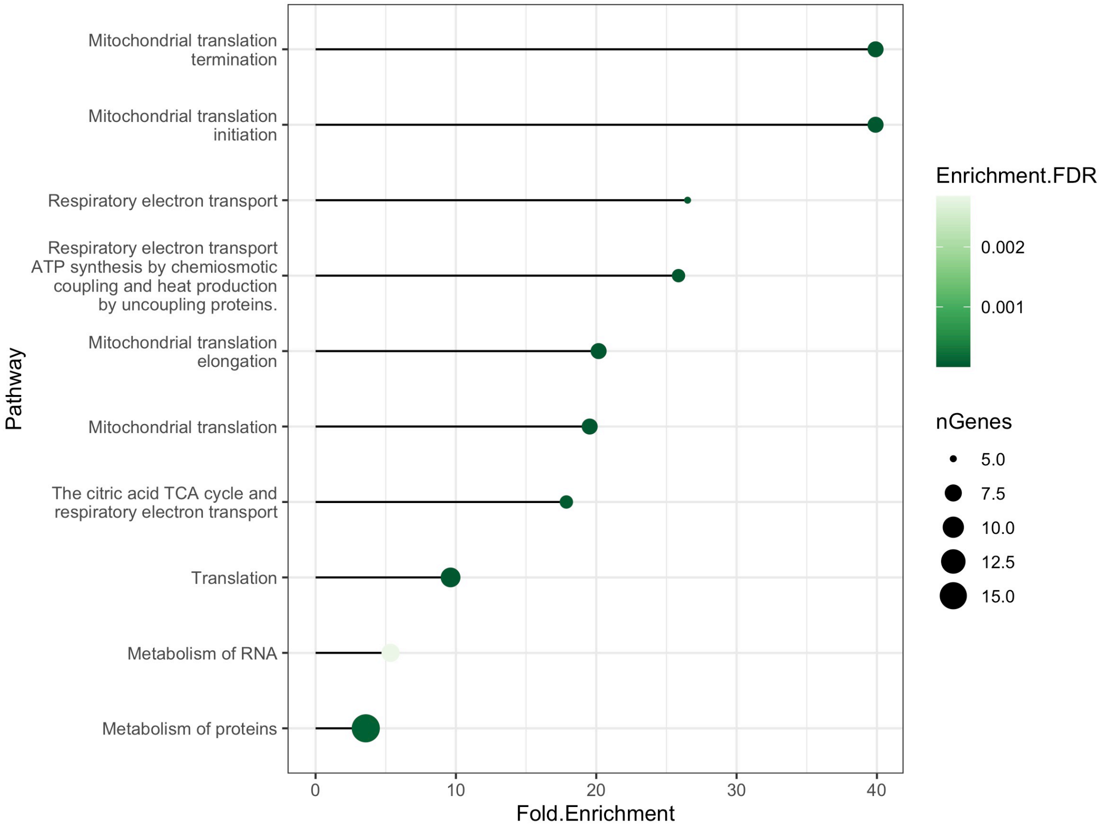


**Fig S.3. Reactome pathway enrichment of lung NET-specific EGs.** The significantly enriched pathways are shown on the y axis; the color bar indicates the significance in terms of False Discovery Rate (FDR)-adjusted p-value, while the dot size indicates the number of genes in the input set found in the pathway. On the x axis the Fold Enrichment, namely the percentage of genes in the input list annotated in a pathway divided by the corresponding percentage in the background human genes.

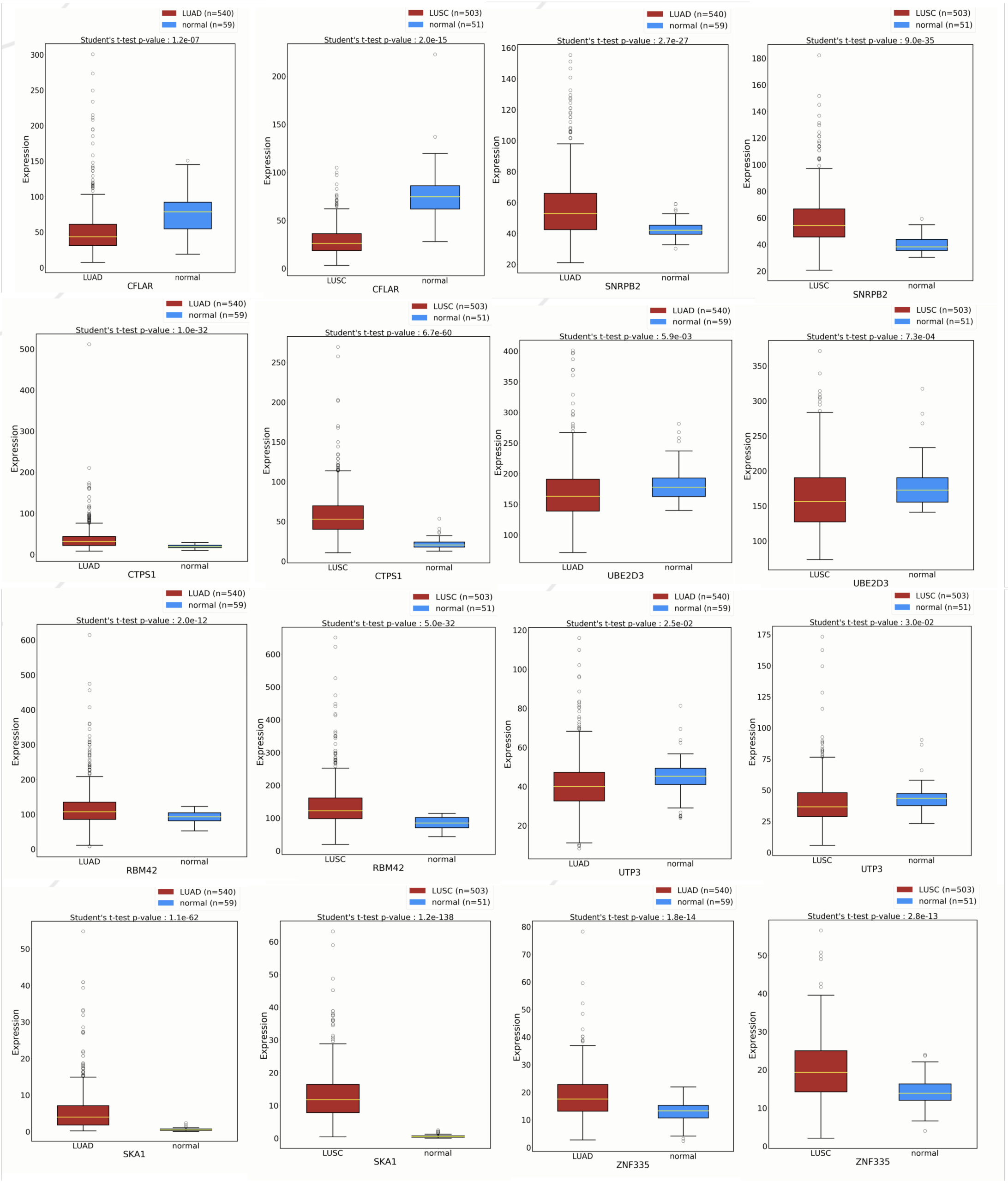


**Fig S.4. Differential expression of NSCLC ucsEGs.** The boxplots show the expression levels of the eight NSCLC-specific EGs in the two NSCLC subtypes, LUAD and LUSC, and normal samples, as collected in OncoDB. The significance of the average difference between the two populations was evaluated with a Student's t-test using the OncoDB platform tool for the differential expression analysis. The legends indicate the colours associated with the groups and the number of samples in brackets.

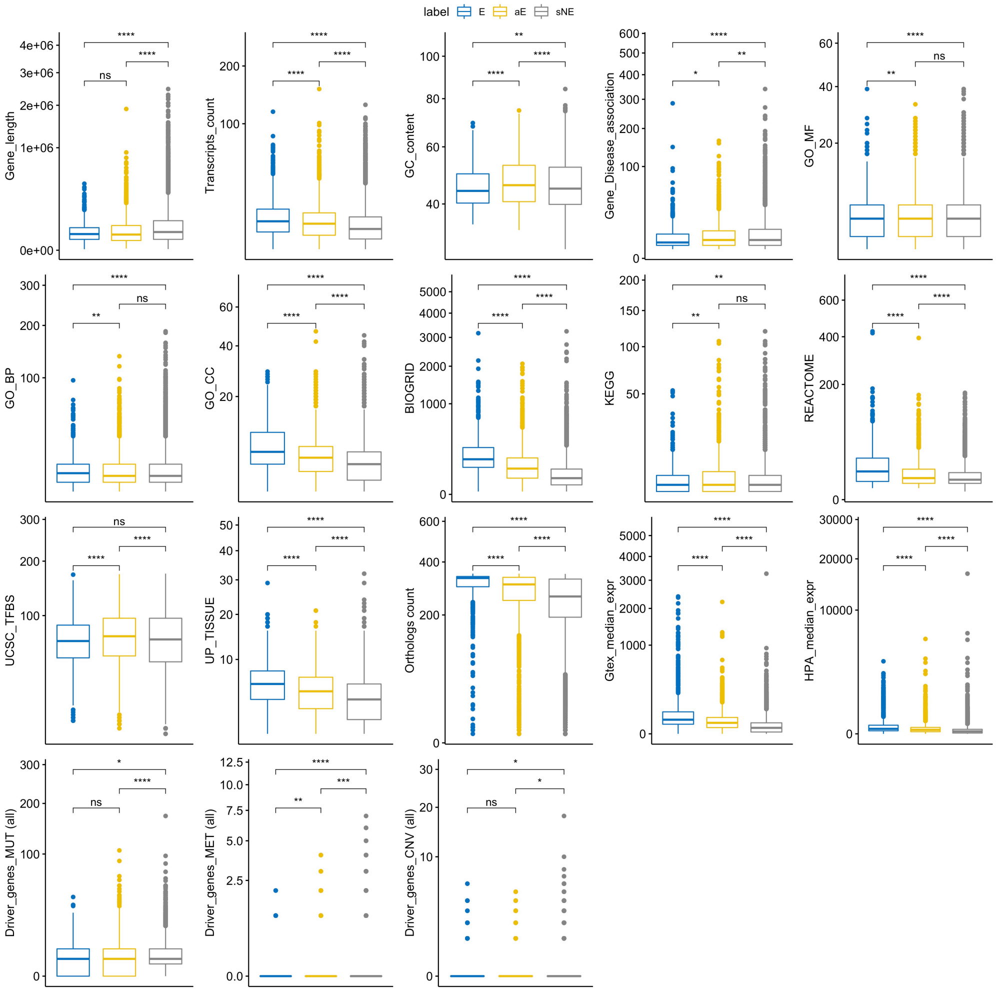


**Fig S.5. Boxplots of the generic Human Bio attribute values for the E, aE, and sNE classes.** The stars on the top indicate the significance of the Wilcoxon test for each pair of comparisons (**** ≤ 0.0001, *** ≤ 0.001, ** ≤ 0.01, * ≤ 0.05, ns = not significant). In favour of visualisation, the values have been signed-square-root transformed.

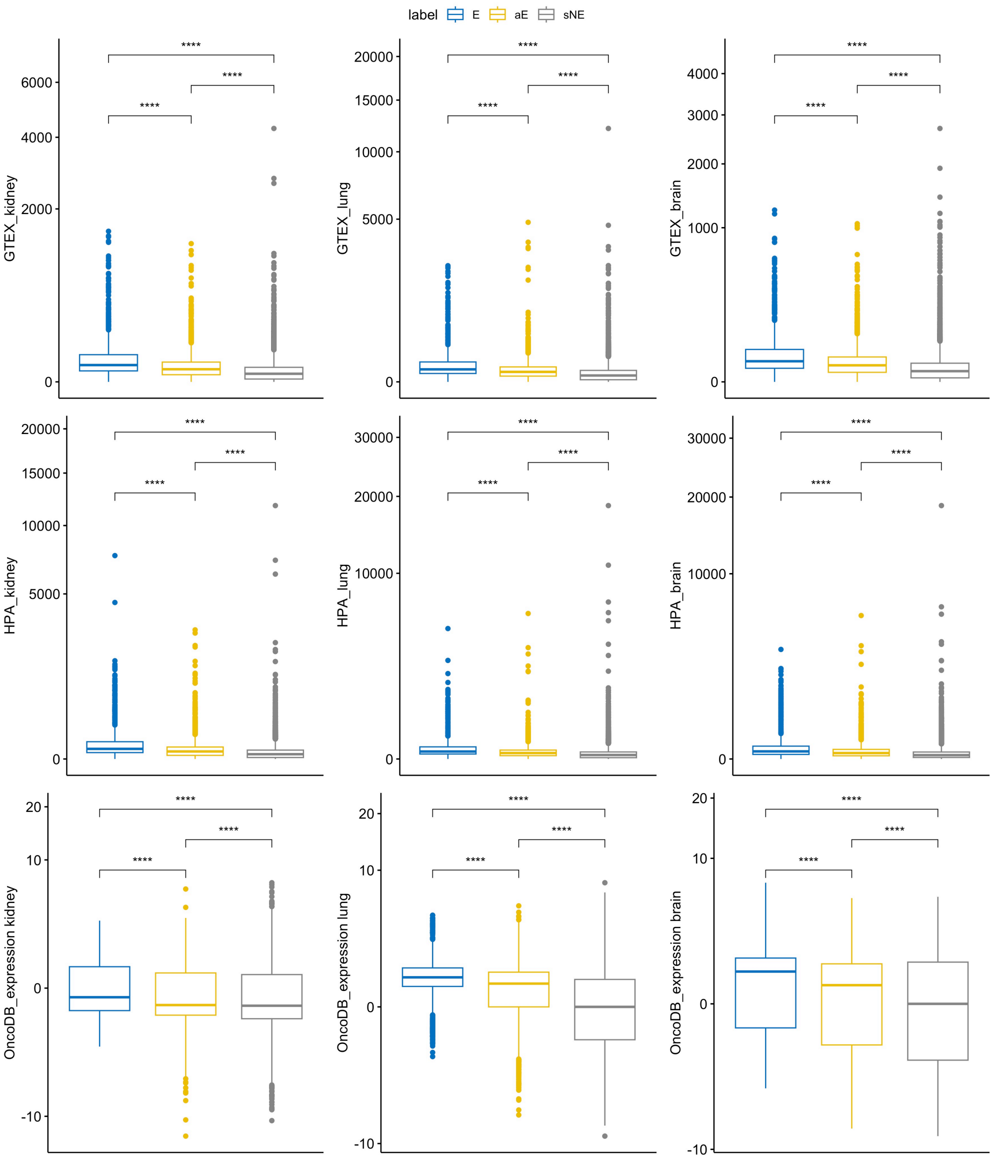


**Fig S.6. Boxplots of the context-specific Bio attribute values of the three tissues investigated for the E, aE, and sNE classes.** The stars on the top indicate the significance of the Wilcoxon test for each pair of comparisons (**** ≤ 0.0001, *** ≤ 0.001, ** ≤ 0.01, * ≤ 0.05, ns = not significant). The Driver genes attributes were not shown as having small ranges of values and poor statistics. In favour of visualisation, the values have been signed-square-root transformed.

(A)

| Generic attribute | ns | * | ** | *** | **** |
|---|---|---|---|---|---|
| Gene length | 96 | 4 | 0 | 0 | aE |
| Transcripts count | 97 | 3 | 0 | 0 | aE |
| GC content | 96 | 2 | 2 | 0 | aE |
| Gene-Disease association | 95 | 5 | aE | 0 | 0 |
| GO-MF | 91;aE | 8 | 1 | 0 | 0 |
| GO-BP | 93;aE | 5 | 2 | 0 | 0 |
| GO-CC | 94 | 4 | 2 | 0 | aE |
| BIOGRID | 94 | 5 | 1 | 0 | aE |
| KEGG | 95;aE | 3 | 2 | 0 | 0 |
| REACTOME | 95 | 5 | 0 | aE | 0 |
| UCSC_TFBS | 93 | 4 | 3 | 0 | aE |
| UP_tissue | 96 | 4 | 0 | 0 | aE |
| Orthologs count | 96 | 2 | 2 | 0 | aE |
| Driver_genes_MUT (all) | 94 | 6 | 0 | aE | 0 |
| Driver_genes_MET (all) | 94 | 4 | 2 | aE | 0 |
| Driver_genes_CNV (all) | 95 | 5;aE | 0 | 0 | 0 |

| cs Attribute | ns | * | ** | *** | **** |
|---|---|---|---|---|---|
| GTEX_kidney | 93 | 4 | 3 | 0 | aE |
| GTEX_lung | 92 | 8 | 0 | 0 | aE |
| GTEX_brain | 94 | 6 | 0 | 0 | aE |
| OncoDB_expression kidney | 94 | 4 | 2 | 0 | aE |
| OncoDB_expression lung | 96 | 2 | 1 | 1 | aE |
| OncoDB_expression brain | 95 | 5 | 0 | 0 | aE |
| HPA_kidney | 95 | 5 | 0 | 0 | aE |
| HPA_lung | 92 | 8 | 0 | 0 | aE |
| HPA_brain | 97 | 1 | 2 | 0 | aE |

(B)

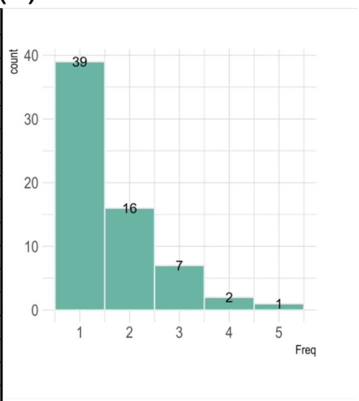


(C)

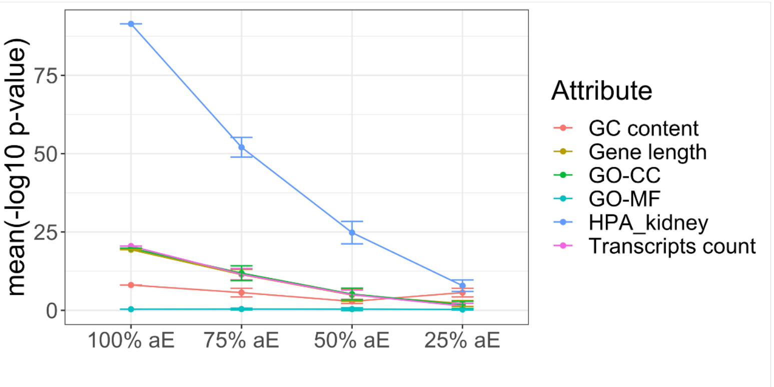


**Fig S.7. Random extraction of the intermediate class.**. A) For each generic attribute (taken as an example from the Kidney dataset) and cs attributes from the three tissues, 100 random partitions of 3000 genes from the sNE groups have been extracted and compared to the rest of the sNE genes. For each tissue, the 100 partitions were fixed. Wilcoxon test was performed to evaluate the statistical significance (p-value) and verify whether the groups come from the same population for each pair of comparisons (**** $\leq 0.0001$, *** $\leq 0.001$, ** $\leq 0.01$, * $\leq 0.05$, ns = not significant). The table indicates the number of partitions for each attribute and for each significance level indicated in the column header. The level of significance given by comparing aE vs sNE, and indicated in Figures S.5;S.6, was also shown by the orange text "aE". B) The histogram shows the number of attributes (x-axis) for which the partitions are simultaneously significant. The count of partitions (y-axis) for each frequency is also shown on the bars. C) The line plot shows the mean of -log10(p-value) and the standard deviation from Wilcoxon tests between different percentages of aE mixed with sNE genes (to 3000 genes) obtained with 10 iterations and the rest of sNE genes for some attributes indicated in the legend.

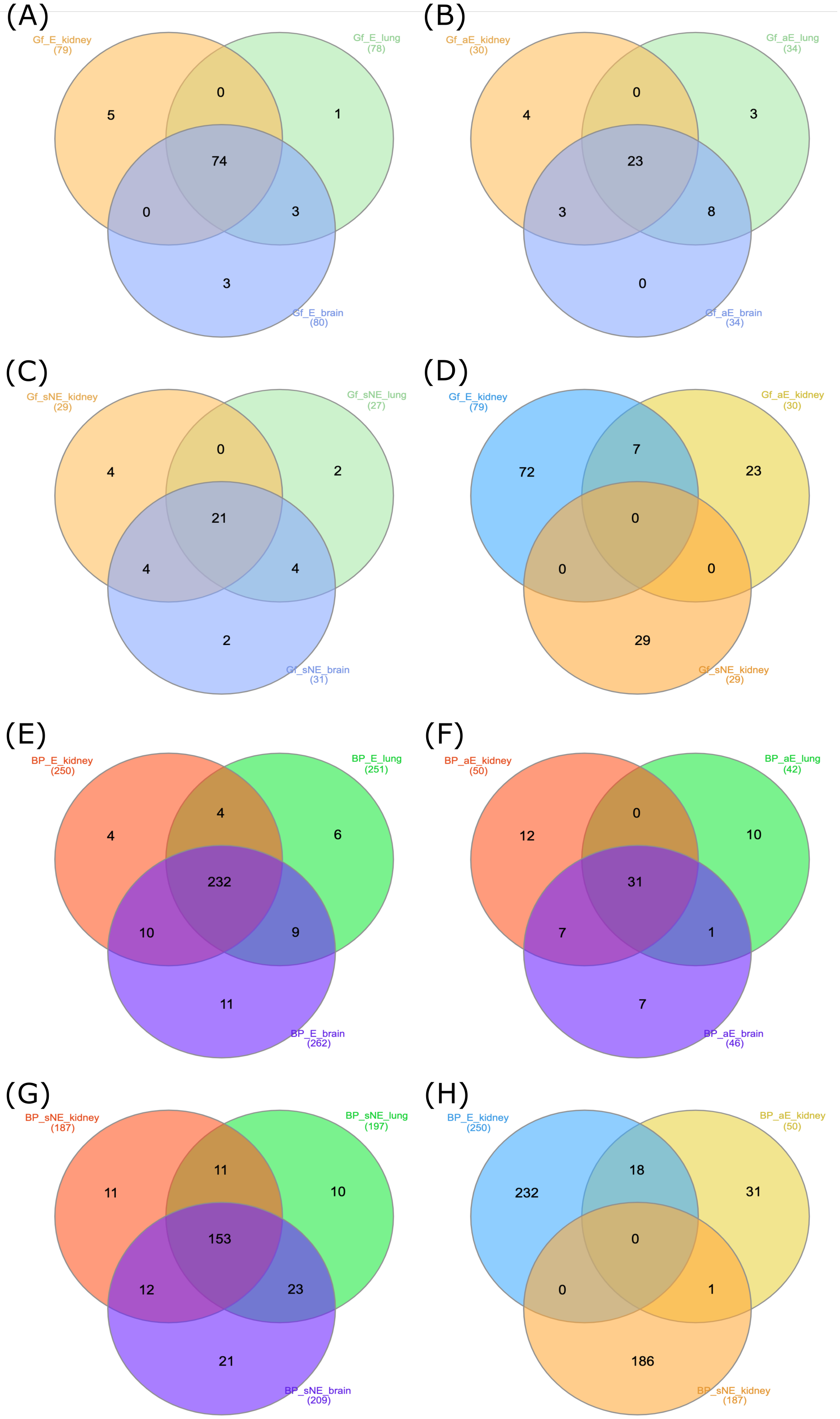


**Fig S.8. Intersection of Gene Families and Biological Processes enrichment among E, aE and sNE genes.** The Venn diagrams show the intersection of Gene Families (gf) and Gene-Ontology Biological Processes (BP) enriched by E, aE or sNE genes among the three tissue contexts under study (A-C; E-G), as well as the intersection of Gene Families (gf) and Gene-Ontology Biological Processes (BP) enriched by genes of the three classes in one context (here Kidney tissue as example) (D;H). The number of genes composing each set is shown in brackets.

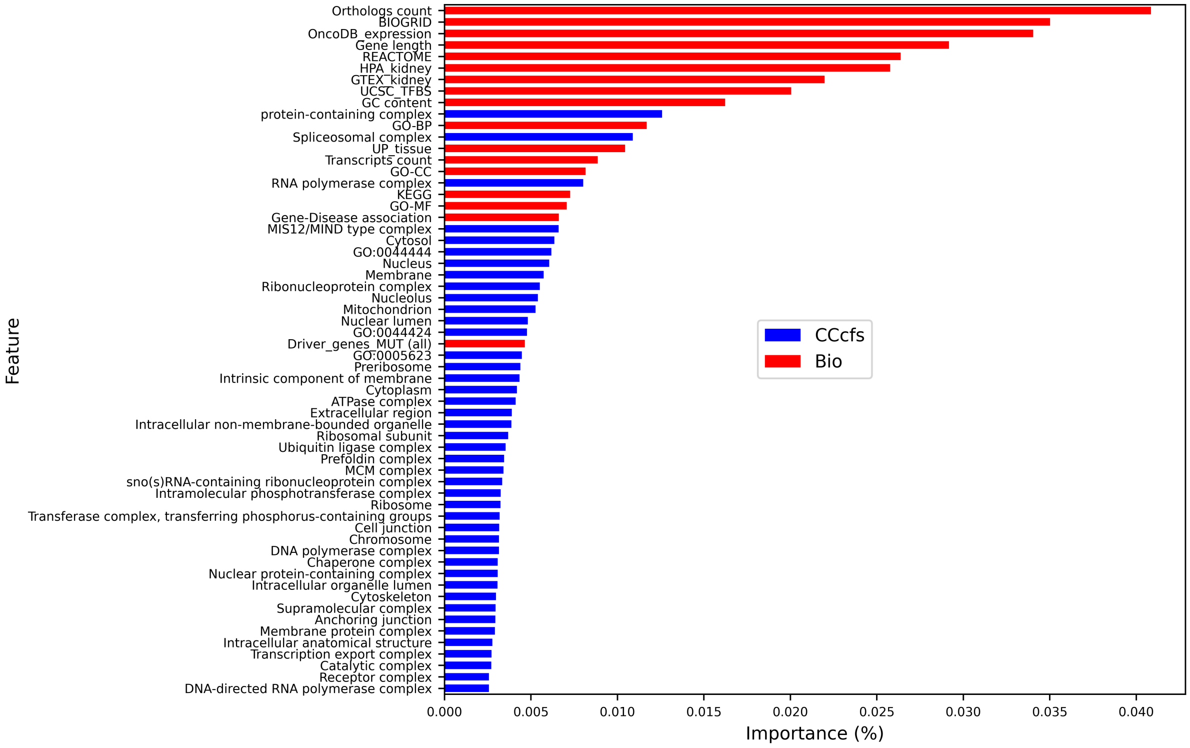


**Fig S.9. Feature importance analysis**. Bio+CCcfs attributes importance calculated by training a sveLGBM model on the entire dataset. The plot cuts-off feature with importance lower than 0.25 %.

### Supplementary Tables

**Table S.1. Collected genomic, transcriptomic, epigenetic, functional and evolutionary features of genes.** (cs) indicates the context-specific attributes.

| Category | Attribute | Description | Data Source |
|---|---|---|---|
| Structure | Gene length | Gene End (bp) - Gene Start (bp) | biomaRt R package v2.54 [51] |
| | GC content | % of Guanosine + Citosine | |
| | Transcripts count | No. of transcripts/gene | |
| Expression | GTEX_* (cs) | Gene median expression in the context of interest | GTEX portal [23] |
| | UP_tissue | Count of annotated expression in tissues | DAVID [24] |
| | OncoDB_expression (cs) | Differential Gene Expression in cancer | OncoDB [25] |
| | HPA_* (cs) | Normalised transcript expression summarised per gene in the context of interest | HPA [26] |
| Function & Localisation | GO-MF | No. of GO-MF annotations | DAVID [24] |
| | GO-BP | No. of GO-BP annotations | |
| | GO-CC | No. of GO-CC annotations | |
| | KEGG | No. of KEGG pathway annotations | |
| | REACTOME | No. of REACTOME pathway annotations | |
| | CCcfs | Subcellular localisation confidence score | COMPARTMENTS [27] |
| Interaction | BIOGRID | No. of BIOGRID interactions annotations | DAVID [24] |
| | UCSC_TFBS | Transcription factors binding sites prediction | |
| Conservation | Orthologs count | No. of orthologous/gene | NCBI [29] |
| Association with Disease | Driver_genes_MUT (cs) | No. of predictions as 'MUT driver' in cancer | DriverDBv3 [30] |
| | Driver_genes_CNV (cs) | No. of predictions as 'CNV driver' in cancer | |
| | Driver_genes_MET (cs) | No. of predictions as 'Methylation driver' in cancer | |
| | Gene-Disease association | No. of associations with diseases | DisGeNet [31] |

**Table S.2. Comparison of classifiers on prediction in "E vs NE" problem in the Kidney case study.** Ranking of methods is based on the Balanced Accuracy metric. All methods with "sve" prefix are our meta-learning model proposal with a different base classifier as member of the ensemble. All other methods are provided by the PyCaret library. All models where trained with Bio+CCcfs+N2V attributes of genes. CPU times are measured on Apple M2 with 16GB RAM.

| Model | Accuracy | ROC-AUC | Sensitivity | Specificity | BA | TT (Sec) |
|---|---|---|---|---|---|---|
| sveLGBM | 0.850100 | 0.951200 | 0.914800 | 0.845000 | 0.879900 | 14.608000 |
| sveADA | 0.856900 | 0.945400 | 0.901100 | 0.853500 | 0.877300 | 13.146000 |
| sveET | 0.866600 | 0.936400 | 0.852700 | 0.867600 | 0.860200 | 3.588000 |
| sveRF | 0.883200 | 0.938600 | 0.832000 | 0.887200 | 0.859600 | 3.008000 |
| Random Forest Classifier | 0.810200 | 0.903600 | 0.830800 | 0.808600 | 0.819700 | 0.916000 |
| Extra Trees Classifier | 0.826100 | 0.871100 | 0.761800 | 0.831100 | 0.796500 | 0.758000 |
| Linear Discriminant Analysis | 0.945500 | 0.931800 | 0.619100 | 0.970900 | 0.795000 | 6.512000 |
| sveLDA | 0.740800 | 0.856100 | 0.837800 | 0.733300 | 0.785500 | 5.074000 |
| Logistic Regression | 0.899400 | 0.842400 | 0.627200 | 0.920500 | 0.773900 | 1.572000 |
| SVM - Linear Kernel | 0.885200 | 0.827900 | 0.600700 | 0.907300 | 0.754000 | 19.138000 |
| Ada Boost Classifier | 0.943700 | 0.928900 | 0.492500 | 0.978700 | 0.735600 | 4.790000 |
| Light Gradient Boosting Machine | 0.947900 | 0.940600 | 0.474100 | 0.984700 | 0.729400 | 2.174000 |

**Table S.3. sveLGBM tuning of parameters with Optuna library**. Optimiziation was carried out on "E vs NE" classification problem with a stratified 5-fold cross-validation with Bio+CCcfs+N2V features by maximising BA metric.

| Trial no. | boosting_type | learning_rate | n_estimators | n_voters | BA |
|---|---|---|---|---|---|
| 37 | gbdt | 0.094505 | 200 | 13 | 0.893151 |
| 15 | gbdt | 0.098300 | 140 | 10 | 0.891459 |
| 44 | gbdt | 0.076452 | 200 | 12 | 0.890954 |
| 43 | gbdt | 0.075168 | 200 | 12 | 0.890826 |
| 41 | gbdt | 0.078591 | 200 | 13 | 0.890602 |
| 33 | gbdt | 0.098020 | 180 | 13 | 0.890241 |
| 31 | gbdt | 0.059095 | 160 | 11 | 0.889936 |
| 34 | gbdt | 0.085756 | 200 | 13 | 0.889739 |
| 22 | gbdt | 0.063759 | 180 | 9 | 0.889298 |
| 30 | gbdt | 0.054934 | 160 | 12 | 0.889146 |
| 36 | gbdt | 0.076796 | 200 | 14 | 0.889028 |
| 23 | gbdt | 0.065602 | 160 | 9 | 0.888994 |
| 42 | gbdt | 0.076634 | 200 | 16 | 0.888759 |
| 40 | gbdt | 0.044127 | 180 | 10 | 0.888419 |
| 4 | gbdt | 0.088891 | 140 | 15 | 0.888175 |
| 39 | gbdt | 0.098960 | 140 | 14 | 0.887998 |
| 49 | gbdt | 0.057674 | 200 | 16 | 0.887826 |
| 11 | gbdt | 0.059871 | 180 | 15 | 0.886745 |
| 47 | gbdt | 0.049777 | 200 | 14 | 0.886566 |
| 29 | gbdt | 0.042902 | 180 | 9 | 0.886259 |
| 32 | gbdt | 0.052158 | 140 | 11 | 0.885557 |
| ... | ... | ... | ... | ... | ... |
| 5 | gbdt | 0.001175 | 100 | 7 | 0.500000 |

**Table S.4. Classification performance metrics adopted in the experiments.** They are defined in terms of the number of true positives (TP), true negatives (TN), false positives (FP), and false negatives (FN), where the first class in each binary task (e.g. class E in the "E vs NE" classification task) is assumed as the positive class.

| **Metric** | **Description** | **Formula** |
|---|---|---|
| Accuracy | % of correctly classified samples | $\frac{TP+TN}{TP+FP+FN+TN}$ |
| Specificity (TNR) | % of negative samples correctly classified | $\frac{TN}{TN+FP}$ |
| Sensitivity (TPR) | % of positive samples correctly classified | $\frac{TP}{TP+FN}$ |
| Balanced Accuracy (BA) | Average of Specificity and Sensitivity | $\frac{1}{2}(\text{Sensitivity} + \text{Specificity})$ |
| ROC-AUC | Area Under the Receiver Operating Characteristic curve | $\int_0^1 \text{Sensitivity}(x)dx$, $x = 1 - \text{Specificity}$ |
| CM | Confusion Matrix | $\begin{array}{c\|c} TN & FP \\ \hline FN & TP \end{array}$ |

Apart from the confusion matrix, all the metrics assume values in [0,1], except ROC-AUC, which ranges in [0.5,1]; higher values indicate better performance.

**Table S.5. "E vs NE" classification performance based on HELP labelling.** (A) Kidney, (B) Lung, (C) Brain tissues, and (D) Human. Averages and errors of metrics are obtained on fifty measurements related to ten times iterated 5-fold cross-validation. The averaged Confusion Matrix (CM) is also shown.

| feature | Bio | N2V | CCcfs | Bio+CCcfs | Bio+CCcfs+N2V |
|---|---|---|---|---|---|
| (A) Kidney | | | | | |
| ROC-AUC | 0.914±0.007 | 0.929±0.008 | 0.940±0.008 | 0.956±0.005 | 0.958±0.006 |
| Accuracy | 0.795±0.007 | 0.845±0.006 | 0.861±0.006 | 0.877±0.005 | 0.880±0.005 |
| BA | 0.832±0.010 | 0.854±0.013 | 0.867±0.012 | 0.887±0.010 | 0.892±0.009 |
| Sensitivity | 0.875±0.020 | 0.864±0.027 | 0.873±0.023 | 0.899±0.020 | 0.905±0.019 |
| Specificity | 0.789±0.007 | 0.843±0.007 | 0.861±0.006 | 0.876±0.005 | 0.878±0.005 |
| CM | pred: NE, E<br>true NE: 12618.1, 3375.9<br>true E: 155.3, 1086.7 | pred: NE, E<br>true NE: 13486.2, 2507.8<br>true E: 169.0, 1073.0 | pred: NE, E<br>true NE: 13763.9, 2230.1<br>true E: 157.7, 1084.3 | pred: NE, E<br>true NE: 14003.7, 1990.3<br>true E: 125.2, 1116.8 | pred: NE, E<br>true NE: 14041.4, 1952.6<br>true E: 117.8, 1124.2 |
| (B) Lung | | | | | |
| ROC-AUC | 0.918±0.006 | 0.931±0.008 | 0.941±0.006 | 0.957±0.005 | 0.959±0.005 |
| Accuracy | 0.800±0.007 | 0.852±0.005 | 0.845±0.014 | 0.878±0.005 | 0.882±0.005 |
| BA | 0.839±0.010 | 0.857±0.011 | 0.864±0.011 | 0.891±0.009 | 0.895±0.009 |
| Sensitivity | 0.884±0.019 | 0.863±0.022 | 0.885±0.031 | 0.905±0.017 | 0.910±0.018 |
| Specificity | 0.793±0.008 | 0.851±0.005 | 0.842±0.017 | 0.876±0.005 | 0.879±0.005 |
| CM | pred: NE, E<br>true NE: 12701.7, 3308.3<br>true E: 142.2, 1081.8 | pred: NE, E<br>true NE: 13619.7, 2390.3<br>true E: 168.2, 1055.8 | pred: NE, E<br>true NE: 13486.1, 2523.9<br>true E: 140.9, 1083.1 | pred: NE, E<br>true NE: 14021.9, 1988.1<br>true E: 116.0, 1108.0 | pred: NE, E<br>true NE: 14078.9, 1931.1<br>true E: 109.7, 1114.3 |
| (C) Brain | | | | | |
| ROC-AUC | 0.916±0.006 | 0.932±0.007 | 0.942±0.007 | 0.958±0.005 | 0.960±0.005 |
| Accuracy | 0.801±0.006 | 0.852±0.007 | 0.847±0.014 | 0.882±0.006 | 0.883±0.006 |
| BA | 0.833±0.008 | 0.859±0.011 | 0.866±0.011 | 0.893±0.008 | 0.895±0.008 |
| Sensitivity | 0.869±0.019 | 0.868±0.024 | 0.888±0.031 | 0.906±0.019 | 0.910±0.018 |
| Specificity | 0.796±0.007 | 0.850±0.008 | 0.844±0.017 | 0.880±0.007 | 0.881±0.007 |
| CM | pred: NE, E<br>true NE: 12747.1, 3262.9<br>true E: 161.4, 1072.6 | pred: NE, E<br>true NE: 13612.7, 2397.3<br>true E: 162.7, 1071.3 | pred: NE, E<br>true NE: 13512.1, 2497.9<br>true E: 137.7, 1096.3 | pred: NE, E<br>true NE: 14094.4, 1915.6<br>true E: 116.2, 1117.8 | pred: NE, E<br>true NE: 14104.2, 1905.8<br>true E: 111.1, 1122.9 |
| (D) Human | | | | | |
| ROC-AUC | 0.909±0.008 | 0.912±0.010 | 0.942±0.008 | 0.957±0.006 | 0.957±0.007 |
| Accuracy | 0.790±0.008 | 0.822±0.007 | 0.843±0.006 | 0.878±0.007 | 0.877±0.007 |
| BA | 0.825±0.011 | 0.831±0.012 | 0.867±0.011 | 0.889±0.011 | 0.888±0.013 |
| Sensitivity | 0.865±0.022 | 0.842±0.023 | 0.896±0.021 | 0.903±0.020 | 0.902±0.023 |
| Specificity | 0.784±0.009 | 0.820±0.007 | 0.839±0.007 | 0.876±0.007 | 0.875±0.007 |
| CM | pred: NE, E<br>true NE: 12541.8, 3450.2<br>true E: 167.7, 1074.3 | pred: NE, E<br>true NE: 13113.1, 2878.9<br>true E: 196.0, 1046.0 | pred: NE, E<br>true NE: 13418.7, 2573.3<br>true E: 129.3, 1112.7 | pred: NE, E<br>true NE: 14003.3, 1988.7<br>true E: 120.8, 1121.2 | pred: NE, E<br>true NE: 13987.9, 2004.1<br>true E: 121.4, 1120.6 |

**Table S.6. Comparison of sveLGBM and CLEARER on OGEE+DEG labelling for the prediction of cEGs.** *Hs Features* refer to the features collected for Homo Sapiens EGs prediction presented in the work [38]. sveLGBM hyperparameters: n_voters=16, learning_rate=0.1, n_estimators=200, boosting_type='gbdt'. CLEARER hyperparameter: RF n_estimators=500 as in [38].

| method | sveLGBM (HELP) | RandomForest (CLEARER) |
|---|---|---|
| metric | Bio+CCcfs+N2V | Hs Features reduced by lasso |
| ROC-AUC | 0.9728±0.0051 | 0.9682±0.0024 |
| Accuracy | 0.9111±0.0068 | 0.9625±0.0025 |
| BA | 0.9130±0.0144 | 0.7844±0.0123 |
| Sensitivity | 0.9152±0.0359 | 0.5834±0.0240 |
| Specificity | 0.9108±0.0090 | 0.9854±0.0019 |
| CM | pred: E, NE<br>true E: 755, 70<br>true NE: 1177, 12019 | pred: E, NE<br>true E: 486, 347<br>true NE: 200, 13543 |

**Table S.7. Comparison of sveLGBM, DeepHE and EPGAT predictions on HELP labelling for Kidney-, Lung-, Brain-specific EGs, and cEGs (Human).** EPGAT running with PPI input and sublocalisation attributes. EPGAT hyper-parameters are optimised by using the provided tuning function. DeepHE running with DNA sequencing extracted features plus node2vec embedding 120-sized features extracted from the PPI. HELP running with Bio+CCcfs + N2V embedding 120-sized features extracted from the PPI.

| | Kidney | | | Lung | | |
|---|---|---|---|---|---|---|
| metric | EPGAT | DeepHE | sveLGBM | EPGAT | DeepHE | sveLGBM |
| AUC | 0.902±0.007 | 0.921±0.016 | 0.957±0.006 | 0.913±0.009 | 0.916±0.021 | 0.958±0.005 |
| Acc. | 0.834±0.028 | 0.845±0.016 | 0.894±0.004 | 0.843±0.032 | 0.845±0.023 | 0.895±0.004 |
| BA | 0.824±0.012 | 0.845±0.016 | 0.890±0.009 | 0.832±0.014 | 0.845±0.023 | 0.892±0.010 |
| Sens. | 0.813±0.045 | 0.866±0.02 | 0.886±0.019 | 0.819±0.051 | 0.877±0.029 | 0.889±0.020 |
| Spec. | 0.835±0.033 | 0.824±0.024 | 0.894±0.004 | 0.845±0.037 | 0.812±0.028 | 0.895±0.005 |
| | Brain | | | Human | | |
| metric | EPGAT | DeepHE | sveLGBM | EPGAT | DeepHE | sveLGBM |
| AUC | 0.908±0.012 | 0.921±0.009 | 0.959±0.005 | 0.880±0.017 | 0.91±0.02 | 0.957±0.007 |
| Acc. | 0.857±0.022 | 0.847±0.012 | 0.898±0.006 | 0.784±0.043 | 0.83±0.027 | 0.891±0.006 |
| BA | 0.833±0.008 | 0.847±0.012 | 0.894±0.009 | 0.798±0.020 | 0.83±0.027 | 0.886±0.013 |
| Sens. | 0.806±0.027 | 0.884±0.022 | 0.890±0.019 | 0.815±0.063 | 0.898±0.037 | 0.880±0.024 |
| Spec. | 0.861±0.026 | 0.811±0.024 | 0.898±0.006 | 0.781±0.050 | 0.762±0.047 | 0.892±0.007 |

**Table S.8. Optimal hyper-parameters of sveLGBM, DeepHE and EPGAT methods used in comparison of Table S.7.**

| method | Kidney | Lung | Brain | Human |
|---|---|---|---|---|
| EPGAT | epochs=1000,<br>lr=0.005,<br>weight_decay=0.0005,<br>h_feats=[8,1],<br>heads=[8,1],<br>dropout=0.4 | epochs=1000,<br>lr=0.005,<br>weight_decay=0.0005,<br>h_feats=[8,1],<br>heads=[8,1],<br>dropout=0.4 | epochs=1000,<br>lr=0.00057,<br>weight_decay=0.000247,<br>h_feats=[32,8, 1],<br>heads=[8,4,1],<br>dropout=0.137 | epochs=1000,<br>lr=0.0023,<br>weight_decay=0.000126,<br>h_feats=[64,1],<br>heads=[4,1],<br>dropout=0.34 |
| DeepHE | epochs=50, batch_size=32, dropout=0.2, h_feats=[128,256,512], folding=1 | | | |
| sveLGBM | n_voters=13, n_estimators=200, boosting_type=gbdt, learning_rate=0.1 | | | |

**Table S.9. "E vs sNE", "E vs aE" and "aE vs sNE" classification performance based on HELP labelling.** The case study is Kidney tissue using Bio+CCcfs+N2V features. Averages and errors of metrics are obtained on fifty measurements related to ten times iterated 5-fold cross-validation. The averaged Confusion Matrix (CM) is also shown.

| problem | E vs sNE | E vs aE | aE vs sNE |
|---|---|---|---|
| ROC-AUC | 0.973±0.004 | 0.895±0.009 | 0.751±0.010 |
| Accuracy | 0.915±0.005 | 0.797±0.012 | 0.713±0.007 |
| BA | 0.915±0.007 | 0.813±0.012 | 0.687±0.010 |
| Sensitivity | 0.916±0.016 | 0.849±0.021 | 0.644±0.019 |
| Specificity | 0.915±0.005 | 0.776±0.016 | 0.729±0.008 |
| CM | pred: sNE, E<br>true sNE: 11790.0, 1096.0<br>true E: 104.8, 1137.2 | pred: aE, E<br>true aE: 2412.3, 695.7<br>true E: 187.7, 1054.3 | pred: sNE, aE<br>true sNE: 9396.4, 3489.6<br>true aE: 1106.2, 2001.8 |
