## Supplementary Methods for "HELP: A computational framework for labelling and predicting human common and context-specific essential genes"

### M.1 Base estimator choice for SVE

In this supplementary section, we report the experience acquired on tuning and comparing, by using the PyCaret toolset [52], the performance of several classifiers on the binary classification problem of E and NE genes, which is characterised by a strong unbalancing of class proportions. The experiment was conducted on the kidney tissue case study. The list of classifiers considered in the comparison includes:

1. classifiers from the PyCaret library: Random Forest Classifier, Extra Tree Classifier, Linear Discriminant Analysis, Logistic Regression, SVM (with linear kernel), AdaBoost Classifier, Light Gradient Boosting Machine (LGBM);

2. our splitting voting ensemble method applied each time with a different base classifier from this list: Random Forest Classifier (sveRF), Extra Tree Classifier (sveET), Linear Discriminant Analysis (sveLDA), AdaBoost Classifier (sveADA), and Light Gradient Boosting Machine (sveLGBM).

Before comparison, all methods were tuned in their parameters using the PyCaret `tune_model` facility, using BA as reference metric. After tuning, the optimised models were applied in a stratified 5-fold cross-validation on the case-study dataset. The performance measurements are reported in S.2 (see `https://github.com/giordamaug/HELP/blob/v2.0/notebooks/compare_models.ipynb` for reproducibility).

### M.2 sveLGBM classifier tuning

To identify the best-performing parameters configuration we performed the hyper-parameters optimisation of sveLGBM by using the Optuna library [53]. The more critical hyper-parameter to set is the number of voters, i.e. the number of classifiers into which the training samples of the majority class are split uniformly. The choice of distributing those samples, during training, among the equal classifiers (voters) of the ensemble has the rationale of solving the strong unbalancing of class labels in the training set. Several other hyper-parameters need to be set, in particular, those configuring each identical LGBM member of the ensemble, such as the type of boosting (default `gbdt`), the learning rate (default 0.1), and the number of estimators (indeed, each LGBM is on its own and ensemble of decision trees). Due to the large number of parameters, we decided to explore the optimisation of sveLGBM by using the Optuna library [53] with boosting type varying in: `gbdt`, `dart`); a learning rate varying from 0.001 to 0.1; a number of voters in the ensemble from 1 to 20 members, and a number of estimators for the LGBM model ranging from 60 to 200. The objective function calculates a 5-fold stratified cross-validation on the input dataset (Bio+CCcfs+N2V attributes), and it maximises the BA metric. The results of the optimisation step are reported in Table S.3 (see `https://github.com/giordamaug/HELP/blob/v2.0/notebooks/optuna.ipynb` for reproducibility).

### M.3 Comparison with CLEARER

CLEARER [38] was designed to predict cellular (CEG) and organismal essential genes (OEG), using the gene labels collected in OGEE [39] and DEG [40] databases. The comparison of sveLGBM with CLEARER predictor was carried out by training both models on those labels with different input data: for the former, we used as input the Bio+CCcfs+N2V features, while for the latter, we used features presented by CLEARER's authors and we applied feature selection by Lasso method, as described in [38]. In the case of CLEARER, we adopted the RandomForest classifier parameters as indicated in the publicly available code available at `https://github.com/ThomasBeder/CLEARER`. We performed a one-shot feature elimination step on the whole input dataset. With the Lasso method, we reduced the feature space size by one order of magnitude (from 41635 to 4067 features). Regarding our feature set, we did not accomplish feature elimination. Indeed, as demonstrated in section M.5, the reduction of the Bio+CCcfs feature set discussed here does not imply performance improvements. For both models we carried out a stratified 5-fold cross-validation on the input dataset (with the same random seed for partitioning). At each validation fold, while sveLGBM was applied on the unbalanced train/test data, in the case of CLEARER, a data pre-processing is done to balance the dataset by using SMOTE [54] resampling method.

It should be noted that due to the intersection of genes with attributes and this label in the OGEE-DEG nomenclatures, in CEG prediction with Hs Features, the distribution of genes was as follows: NE=13743, E=833. In the prediction with Bio+CCcfs+N2V features, the distribution was NE=13196, E=825.

For obvious reasons of copyright of the CLEARER software, we did not consider it appropriate to make versions of this code adapted for the purposes of this paper available in public repositories.

### M.4 Comparison with DeepHE and EPGAT

In this supplementary section, we report the performance of methods sveLGBM, DeepHE and EPGAT in predicting E/NE genes in human and three tissue-specific case studies.

We built each prediction model using as input gene attributes the ones described in the reference papers: in particular, DeepHE exploits DNA sequence features, here called "seq" attributes, in combination with node2vec-based embedding of PPI network, here called "embed" features (for addressing tissue-specific and human genes prediction we considered tissue-specific and human PPI, respectively). It should be noted that the latter feature set is the same as the one used to build our prediction model, while we also added "Bio" and "CCcfs" information to the information extracted from PPI, as described in paragraph Features' sets. For EPGAT, we found out by experiments that the best attribute input in all tissue case studies consisted of the sublocalisation attribute set, as also the authors stated in their work [7], which was processed during GAT training according to the topology of the input tissue-specific PPI. Note how, in the case of EPGAT method, the PPI information is not pre-calculated into embedding vectors to be fed to classifiers, whereas it is a permanent adjacency matrix of nodes in the layers of the GAT neural network.

Hyper-parameter optimisation of methods was conducted in the following manner: for EPGAT we used the `hyper_search` functions provided by authors in the public software, while in the case of sveLGBM we used the same optimal hyper-parameters found with Optuna [53] library in the experiments of S.5 which are also reported in Table S.3. DeepHE software does not provide any hyper-parameter optimisation facility, and it uses a fixed configuration of parameters that we suppose is considered optimal by authors; we only varied the `folding` parameters, which regulates the amount of undersampling of majority class samples: in particular to force the DNN underlying model behaving with a higher sensitivity we found out that a 1:1 folding proportion of NE:E samples was the best choice. For obvious reasons of copyright of the DeepHE and EPGAT software, we did not consider it appropriate to make versions of those code adapted for the purposes of this paper available in public repositories.

### M.5 Feature importance analysis

We investigated the importance of features in the context of E versus NE genes classification. The experiment was conducted on kidney-tissue context. The results of this study are reproducible in the notebook `feature_importance.ipynb` on the GitHub software distribution.

To this aim, we used the Bio+CCcfs set of attributes: we decided to skip N2V embedding features since these attributes are automatically extracted by deep learning and the embedding size was already chosen to optimise the performance of a predictor built upon solely embedding features. In addition, we preferred to evaluate feature importance ranks when considering Bio and CCcfs set jointly. Importance ranks of attributes are normalized such that they all sum up to the unity (100% of contribution). Although the two sets have cardinalities different in two orders of magnitudes, we expected that several Bio attributes have larger importance in the classification than single CCcfs attributes.

We used the optimal hyper-parameters for sveLGBM. We cut off the attributes with an importance rank lower than 0.25 %. Nine Bio attributes were the top-most important features (Fig S.9). Except the `Driver_genes_MUT (all)` attribute, all the other `Driver_genes_*` features were less significant (less than

0.25 %). From the importance plot we derived that the global contribution of Bio attributes is 31%, while the remaining 69% of the contribution was due to the sum of the large number of CCcfs features. The feature importance analysis here discussed is reproducible by executing the `https://github.com/giordamaug/HELP/feature-importance.ipynb` notebook.

We exploited the results of this analysis to reduce the large number of CCcfs features (3305) to those having ranks greater than 0.001. With this feature reduction (17 Bio + 167 CCcfs attributes), we conducted a 5-fold cross-validation and evaluated the performance, noticing that the average BA (over ten iterations of the experiments) degraded by 1-2 %.
